# Structural engraftment and topographic spacing of transplanted human stem cell-derived retinal ganglion cells

**DOI:** 10.1101/2020.07.14.196055

**Authors:** Kevin Y Zhang, Caitlyn Tuffy, Joseph L Mertz, Sarah Quillen, Laurence Wechsler, Harry A Quigley, Donald J Zack, Thomas V Johnson

## Abstract

Retinal ganglion cell (RGC) replacement and optic nerve regeneration hold potential for restoring vision lost to optic neuropathy. Following transplantation, RGCs must integrate into the neuroretinal circuitry in order to receive afferent visual signals for processing and transmission to central targets. To date, the efficiency of RGC retinal integration following transplantation has been limited. We sought to characterize spontaneous interactions between transplanted human embryonic stem cell-derived RGCs and the recipient mature mammalian retina, and to identify and overcome barriers to the structural integration of transplanted neurons. Using an *in vitro* model system, following transplantation directly onto the inner surface of organotypic mouse retinal explants, human RGC somas form compact clusters and extend bundled neurites that remain superficial to the neural retinal tissue, hindering any potential for afferent synaptogenesis. To enhance integration, we explored methods to increase the cellular permeability of the internal limiting membrane (ILM). Digestion of extracellular matrix components using proteolytic enzymes was titrated to achieve disruption of the ILM while minimizing retinal toxicity and preserving endogenous retinal glial reactivity. Such ILM disruption is associated with dispersion rather than clustering of transplanted RGC bodies and neurites, and with a marked increase in transplanted RGC neurite extension into retinal parenchyma. The ILM appears to be a barrier to afferent retinal connectivity by transplanted RGCs and its circumvention may be necessary for successful functional RGC replacement through transplantation.

## Background

Retinal ganglion cell (RGC) replacement provides a possible therapeutic strategy to reverse vision loss from optic neuropathies such as glaucoma, the world’s leading cause of irreversible blindness.^1,2^ Promising photoreceptor transplantation studies^3-6^ (including human-rodent xenografts^7–11^) provide proof of principle that vision restoration may be attainable by mammalian retinal cell replacement. However, unlike photoreceptors, RGCs are projection neurons and their functional replacement requires bidirectional visual pathway integration. Studies of endogenous RGC axon regeneration following injury identify numerous molecular pathways that can be targeted to drive optic nerve regeneration and efferent visual signal propagation by exogenous transplanted RGCs.^12–17^ However, factors limiting transplanted RGC somal migration, spatial patterning, or dendrite integration within the recipient mammalian retina, all of which are necessary for achieving afferent input, are equally important to functional vision restoration and remain comparatively understudied.

A number of groups have transplanted neural-progenitors and RGC precursor cells into rodent eyes with varying degrees of survival, but clear evidence of functional RGC replacement has been elusive.^18–27^ Human ES-derived RGCs survive in rat eyes and may localize to the RGC layer (RGCL), though these cells did not extend neurites and were RBPMS-negative.^28^ Recently, a pivotal study yielded qualified success in transplanting primary mouse RGCs into rat recipients, documenting relatively rare instances of mature RGC morphology, structural synaptogenesis, and functional electrophysiologic responses to light.^29^ Although encouraging, progression from these initial studies to functional and clinically relevant RGC transplantation requires significantly increasing the efficiency of retinal integration of transplanted cells, a goal that would be aided by better understanding the key barriers that impede integration.

Gene therapy studies show that the internal limiting membrane (ILM) is a major barrier to retinal neuronal transduction by intravitreally administered viral vectors.^30–33^ The ILM is a basement membrane composed of extracellular matrix (ECM) proteins including laminin, collagen IV, perlecan, nidogen, and others.^34^ Cellular interactions with the ILM play important developmental roles in retinal patterning of neurons, glia, and blood vessels.^35–38^ Some have speculated that the ILM could impede retinal integration of various cell types following intravitreal transplantation.^39–43^ We directly evaluated ILM’s impact on intraretinal migration of mesenchymal stem cells (MSCs) specifically and showed that reactive gliosis, not the ILM, impedes engraftment.^44^ However, the role of the ILM in engraftment of transplanted neurons, including RGCs, has not been directly investigated.

During development, spatial localization of RGCs into a tiled mosaic is important for retinotopic patterning. RGC localization ultimately results from coordinated cell differentiation, migration, and selective apoptosis to achieve non-overlapping dendritic fields of similar RGC subtypes.^35,45–48^ RGC interactions with the ILM are important for normal spatial patterning.^35,38^ Given modest success of RGC transplantation thus far, it remains unclear whether and how transplanted RGCs might spread to cover the retina. Characterizing spatial localization patterns of transplanted RGCs in a quantitative manner is key to developing methods for ensuring coverage that recapitulates functional retinotopic maps.

Since rodent and primate RGC physiology are driven by fundamentally divergent gene expression profiles,^49^ studying human RGC transplantation is critical to clinical translation. Several laboratories have developed methods for generating RGCs from human stem cells.^50–52^ Previously, we genome-engineered human embryonic stem (hES) cells to expresses fluorescent reporters under control of the *BRN3B* gene. We optimized a soluble factor-based differentiation protocol to efficiently produce and immunopurify RGCs, and we reported their transcriptomic and electrophysiological characteristics (henceforth referred to as hES-RGCs).^52–54^

Herein, we examine the survival and morphology of hES-RGCs following transplantation onto adult murine organotypic retinal explants to characterize their potential for spontaneous retinal engraftment. We chose the retinal explant model based on extensive prior characterization demonstrating tissue viability for 14 days with progressive endogenous RGCs death that models optic neuropathy.^44,55–57^ Retinal explants exclude MSCs from engraftment in a manner similar to that observed after injection into the living eye.^44^ Using this model we demonstrate that the ILM plays an important role in hES-RGC topographic spacing and neurite localization within the retinal parenchyma.

## Materials and Methods

### Animals

Adult (age 8-16 weeks) CD1 mice or C57BL/6-Tg(CAG-EGFP)10sb/J mice that express GFP ubiquitously (Jackson Laboratories, Bar Harbor, ME) of both sexes were used. Animals were housed in environmentally controlled (12-hour light/dark cycle), conditions with food and water available *ad libitum.* All experimental procedures were approved by Johns Hopkins University’s Animal Care and Use Committee.

### Human stem cell derived RGCs

Human H9 ES cells (WiCell, Madison, WI) carrying genes for tdTomato and the murine cell-surface protein CD90.2/Thy1.2 driven by the endogenous *POU4F2* (*BRN3B)* promoter were clonally propagated in mTeSR-1 media (StemCell Technologies, Cambridge, MA) on growth factor-reduced Matrigel substrate (Corning, Corning, NY), in 10% CO_2_/5% O_2_. Differentiation to RGC fate and immunopurification were performed as described previously.^52^ See Supplemental Methods.

### Organotypic retinal explants

Neural retina was separated from the retinal pigmented epithelium and flat mounted for culture on polytetrafluoroethylene organotypic filters (Millipore-Sigma, Burlington, MA) with the photoreceptor side against the membrane, as described previously.^44,55–57^ See Supplemental Methods.

### Proteolytic enzymes

Proteolytic enzymes in 5μL aliquots were applied to the inner (vitreous) surface of organotypic retinal explants and incubated for 30-60min at 37°C, inactivated by bathing explants in ovomucoid (10mg/mL, Millipore-Sigma) and bovine serum albumin (BSA, 10mg/mL, Millipore-Sigma) in BSS for 5min at 37°C, washed twice in phosphate-buffered saline (PBS), and placed back into culture media. RGC transplantation occurred ≥24h later.

### Light Microscopy

All samples were reassigned random identification numbers by a second investigator to mask the microscopist and ensure unbiased field selection for imaging. Unmasking occurred only after microscopy and image analysis. Cryosections and retinal explant flat-mounts were imaged using confocal laser scanning microscopes (Model 510 or 710, Carl Zeiss Microscopy, Thornwood, NY). Images were obtained with a Plan-Apochromat 40x/1.3 Oil DIC M27 objective, measured 212.34μm x 212.34μm (x,y), and were acquired with voxel size 0.208μm x 0.208μm x 0.449μm (x,y,z). The pinhole was set to 1 Airy unit. Random fields of retinal explant flat-mounts were selected for microscopy, but areas of retina within 300μm of the tissue edge, relaxing incisions or any obvious tissue trauma were excluded.

Cryosections and flat-mounts underwent epifluorescent imaging using an EVOS microscope (Life Technologies, Grand Island, NY). Individual fields were manually focused and imaged using the 20x objective. Image tiles were stitched to form one single image per sample. Retinal explant cryosections and retinal flatmounts were imaged and analyzed in their entirety (i.e. not sampled).

### Image analyses

Topographic localization of RGCs on retinal explants was analyzed using ImageJ (v1.52u, NIH, Bethesda, MD). Nearest neighbor (NN) distance was determined using a script developed for ImageJ.^58^ Density recovery profiles (DRPs) were generated using the sjedrp R package (Stephen Eglen, University of Cambridge).^59^ L(r)-r derivatives of Ripley’s *K*-function were generated using the Lest function from the R spatstat package.

Analyses of hES-RGC neurites were performed using Imaris (v9.3, Oxford Instruments, Zurich, Switzerland). To parse neurite localization according to retinal layer, 3D reconstructions were visualized and rectangular surfaces were created manually spanning the x-y dimensions of the image volume, with z dimensions corresponding to retinal layer boundaries visualized by DAPI stain, which included: a superficial layer that included cells external to the retina and within the retinal nerve fiber layer (RNFL) and RGCL; inner plexiform layer (IPL); inner nuclear layer (INL); and outer nuclear layer (ONL). Resolving the outer plexiform layer OPL or the RGCL separately from the overlying superficial layer of transplanted cells was attempted and inconsistent across entire 3D reconstructed volumes due to tissue undulation. hES-RGC neurites superficial to the retina, within the RNFL, or within the RGCL would all be incapable of synapsing within afferent retinal neurons in the IPL, and therefore were treated similarly. The tdTomato signal was masked according to retinal layer surface. Neurites were traced in a semi-manual manner using the filament workflow and the autopath tool.

hES-RGC soma localization within recipient retina was assessed in Zen software (v8.1.0, 2012 SP1, Carl Zeiss Microscopy) using z-stack scrolling and orthogonal projections. Co-localization between tdTomato and GFP was evaluated in Zen using co-localization histograms and orthogonal projection. Co-localization between tdTomato and PSD-95 puncta was evaluated in Imaris using the co-localization tool.

### Statistical analyses

Data are reported as mean±standard deviation unless otherwise stated. At least 4-6 separate biological samples were analyzed per group. Each experiment was performed at least twice with independent biological replicates. Group means were compared using unpaired two-tailed t-tests or one-way analyses of variance (ANOVA), and pairwise comparisons were made with Dunnett’s post-hoc tests with corrections for multiple comparisons to a single control group. *χ^2^* tests were used to compare the frequencies of gene expression within hES-RGCs according to retinal localization. P<0.05 following correction for multiple comparisons was considered statistically significant. Data were analyzed using SPSS (v25, IBM Corp, Armonk, NY) and plotted using Prism (v8.0, GraphPad Software, San Diego, CA). The data that support the findings of this study are available from the corresponding author upon reasonable request.

## Results

### Survival and topographic localization of transplanted hES-RGCs

We transplanted hES-RGCs in a 5μL single cell suspension at three doses (1.3×10^4^, 2.5×10^4^, or 5.0×10^4^ cells/retina) onto the inner surface of adult mouse organotypic retinal explants. Following 1 week of co-culture, 12.6±8.2% of transplanted hES-RGCs survived (average of the 3 doses). The lowest transplantation dose exhibited the lowest survival rate (Fig 1A). Microscopic evaluation of retinal tissue as a flatmount permitted examination of the two-dimensional spatial arrangement of surviving hES-RGCs and their neurites (Fig 1B-F). Predominantly, hES-RGC somas concentrated within clusters with direct contact between adjacent cell bodies. Outside of clusters were sizeable spaces devoid of hES-RGC somas. We identified relatively few dispersed single cells. Individual neurites and compacted linear neurite bundles extended from cell clusters (on average 6.8 bundles/100 hES-RGCs). Neurites possessed terminal structures resembling growth cones.

**Figure 1.**
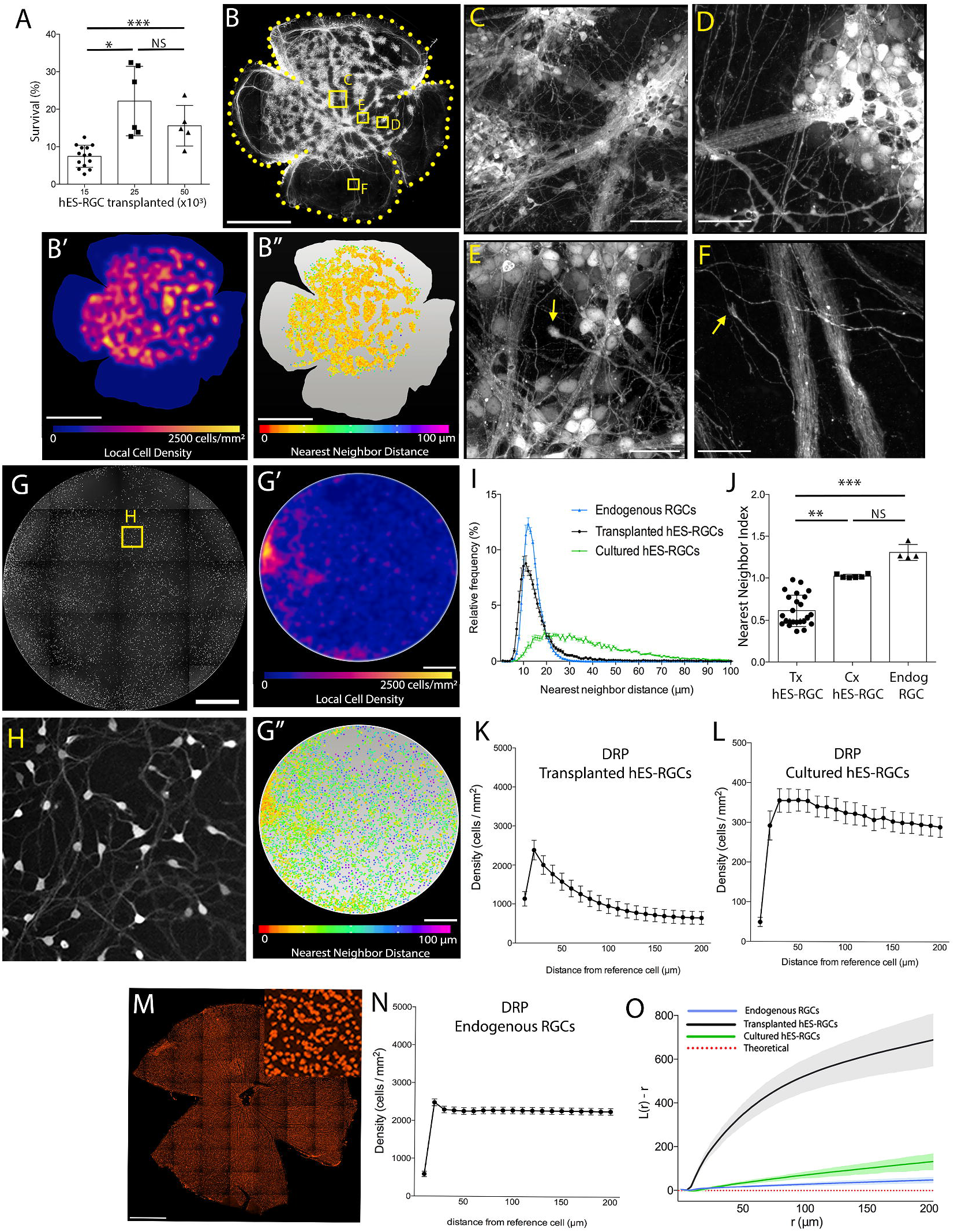
Topographic spacing of hES-RGCs in following transplantation onto neural retina. Human embryonic stem cell (hES) derived retinal ganglion cells (RGCs) were transplanted (Tx) onto the surface of organotypic retinal explants or cultured (Cx) on poly-L-ornithine and laminin-coated for 1 week. RGC survival was lower when fewer cells were transplanted (A). Epifluorescence microscopy revealed the morphology and spacing of tdTomato^+^ hES-RGCs cultured on retinal explants (B-F) vs on laminin-coated polystyrene (G, H). Arrows point to growth cone-like structures. Heat maps (B’, G’) show local cell density and nearest neighbor distance (NND) maps (B’’, G’’) show the distance between each cell and his nearest neighbor, which is also plotted as a histogram (I). Nearest neighbor index normalizes the mean NND to conditions of complete spatial randomness (CSR); values <1.0 indicate clustering. Density recovery profiles (DRP, K, L, N) demonstrate the mean RGC density as a function of distance from each RGC in the sample. Ripley’s L function (O) normalizes the DRP to theoretical CSR conditions such that positive deviations indicate clustering (shaded areas indicate 95% confidence interval). Comparisons to RBPMS-expressing endogenous RGCs in adult C57BL/6 mouse retina are shown (M). Scalebars: 1.25mm (B, G, M); 50μm (C-F). Error bars: standard deviation (A, J); standard error of the mean (I, K, L, N). *p≤0.05, **p≤0.01, ***P≤0.001, NS: p>0.05.

We quantified the topographic spatial clustering of hES-RGCs on retinal explants and compared this to endogenous RGCs immunolabeled for RBPMS (Fig 1M). The overall average density of transplanted hES-RGCs was 253.9±243.3 cell/mm^2^ covering 38.8±11.0% of the retinal surface area. The hES-RGC density within clusters, however, was 2587.5±1394.2 cell/mm^2^, similar to the overall density of endogenous RGCs (2332.8±263.7 cell/mm^2^). The average transplanted hES-RGC nearest neighbor (NN) distance was 16.8±4.5 μm (Fig 1I), similar to what we and others^60,61^ measured for endogenous RGCs (13.5±0.5 μm, Fig 1I). The NN index (NNI) normalizes the NN distance to theoretical conditions of complete spatial randomness (CSR) and measured 0.6±0.2 in transplanted hES-RGCs, indicating cell clustering given a value <1.^62^ By comparison, NNI was 1.3±0.1 for endogenous RGCs (Fig 1J). DRPs, representing the average local spatial density of neighboring RGCs as a function of distance from each index RGC, demonstrated an expected peak followed by plateau for endogenous RGCs (Fig 1N), indicating spatial regularity. In contrast, hES-RGC DRPs demonstrated a rapid exponential decline following the peak (Fig 1K), indicating cell clustering. L(r)-r describes the DRP’s deviation from CSR to objectively compare spatial clustering between experimental groups, which is indicated by a positive deviation from zero. L(r)-r for transplanted hES-RGCs rose steeply over CSR at short distances, whereas endogenous RGCs were only modestly more clustered than CSR (Fig 1O). Across multiple experiments, we transplanted freshly isolated hES-RGCs or thawed cryopreserved hES-RGCs and noted no systematic differences in cell survival or clustering when transplanted onto retinal explants (data not shown).

For comparison, we evaluated spacing of freshly isolated hES-RGCs plated at 250 cells/mm^2^ on poly-L-ornithine and laminin-coated polystyrene. After 1 week, hES-RGC survival was significantly greater than when cultured on retinal explants (81.6%±12.7%, p<0.001). hES-RGCs in cell culture elaborated long neurite processes intertwined in a complex lattice, but we identified no compact neurite bundles (Fig 1G,H). Cell somas were evenly dispersed, with only occasional clusters at the well periphery and very little inter-somal contact (Fig 1G). The average NN distance was 38.5±2.6 μm (Fig 1I). The NNI was 1.03±0.02, indicative of CSR (Fig 1J). The DRP demonstrated a modest, gradual decline after peaking at 30um, indicating minimal clustering (Fig 1L) and L(r)-r showed significantly less clustering than when hES-RGCs were transplanted onto retinal explants (Fig 1O). We performed a similar analysis using thawed cryopreserved hES-RGCs (Supp Fig S1A), which similarly failed to demonstrate significant soma or neurite clustering, but rather exhibited metrics consistent with spatial regularity (Supp Fig 1B,C).

In sum, hES-RGCs exhibited lower survival, compact bundling of long neurites, and clustering of hES-RGC somas when cultured on organotypic retinal explants as compared to poly-L-ornithine and laminin-coated polystyrene. Therefore, local intrinsic retinal factors may impair survival and induce cell body and neurite clustering after transplantation.

### Spontaneous structural engraftment of transplanted hES-RGC neurites

We next examined the 3D structural arrangement of transplanted RGCs and their neurites in relation to the recipient neuroretina. Tracking individual processes within the neurite network extending from transplanted hES-RGCs clusters was not feasible in sectioned tissue because neurites entered and exited the plane of section. We therefore acquired confocal microscopic z-stacks of retinal flat-mounts for volumetric analysis (Fig 2A,D). Away from cut tissue edges, hES-RGC localization within the recipient retina was negligible – nearly all hES-RGC somas and neurites remained in a transplanted cell layer distinctly superficial to the RGCL (Fig 2A, Video 1). Cryosections confirmed that, away from cut edges, hES-RGC somas and neurites were not present within the retinal parenchyma (Fig 2B,C).

**Figure 2.**
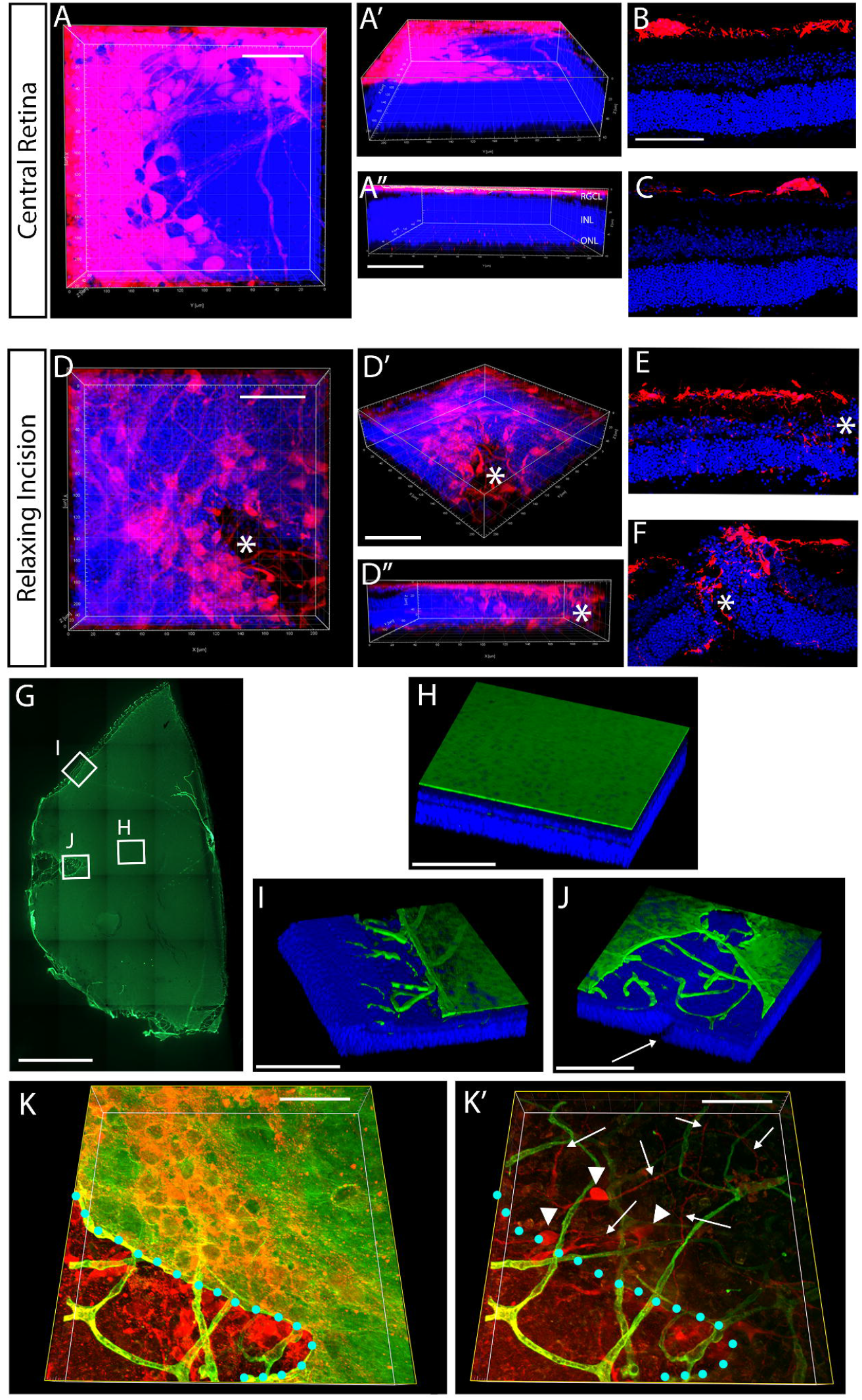
Structural ingrowth of transplanted retinal ganglion cells is greater near sites of physical retinal and internal limiting membrane disruption. Three dimensional reconstructions of confocal microscopy z-stacks are shown (A, D, G-K). Transplanted human embryonic stem cell (hES) derived retinal ganglion cells (RGCs, red) extended neurites which remained superficial to the neural retina when located centrally in the retinal explant (A-C). Near retinal explant edges or sites of relaxing incisions, hES-RGC cell bodies and neurites migrated into the neural retina (D-F). Cryosections from separate retinal explants are shown (B, C, E, F). *Site of retinal disruption (D-F). Immunohistochemistry for laminin (green) shows intact and continuous internal limiting membrane (ILM) in the central retina, with discontinuity at the edges and at relaxing incisions (G-J). Nuclei are labeled with DAPI (blue). Higher magnification images (H-J) demonstrate that underlying retina and retinal vasculature are exposed at peripheral areas of mechanical ILM disruption. J: arrow highlights retinal discontinuity caused by a relaxing incision. Near areas of ILM discontinuity, hES-RGC cell bodies migrate laterally underneath the ILM and extend neurites laterally through the retinal tissue. K and K’ show a 3D reconstructed block of retinal tissue viewed from the top down. K’ is the same block with the most superficial confocal slices that include the ILM removed to reveal the underlying vasculature and hES-RGCs. The edge of the ILM is marked in teal dots. Arrowheads point to hES-RGC somas and arrows point to hES-RGC neurites. RGCL, retinal ganglion cell layer; INL, inner nuclear layer; ONL, outer nuclear layer. Scalebars: 1.25mm (G); 50μm (A,B,D,H-K).

In contrast, near cut edges of the retinal explant, hES-RGCs infiltrated all retinal layers (Fig 2D-F, Video 2). Precise quantification of hES-RGC localization to individual retinal layers was not feasible due to local disruption of the laminar architecture near cut edges. Intraretinal hES-RGC soma and neurite density was greatest near the edges and decreased with distance from the edge. This pattern was consistent with neurite entry from the retinal tissue edge and lateral intraretinal migration, which suggested that a barrier to hES-RGC retinal ingrowth exists at a location superficial to the RGCL, where the ILM is positioned.

We hypothesized that the ILM obstructs structural ingress of transplanted hES-RGCs. Examination of flat-mounted retinal explants confirmed homogenous ILM integrity in central flat-mounts. However, near cut edges the ILM was abruptly broken and neural retina containing laminin^+^ vasculature was exposed deep to retracted ILM (Fig 2G-J). Indeed, hES-RGC somas and neurites were identified deep to the ILM only in areas directly adjacent to retinal explant edges where the ILM was mechanically disrupted (Fig 2K).

### ILM disruption by enzymatic digestion

In order to increase cellular permeability of the ILM, we evaluated enzymatic methods of ECM digestion in retinal explants. We sought to identify an approach that strongly disrupts structural proteins at the ILM while minimizing off-target toxicity to inner retinal neurons and glia, so that the effect of enzymes could be attributed specifically to its effects on the ECM.

We tested several proteolytic enzymes, applied at multiple concentrations directly onto the retinal explant ILM surface and then inactivated them with ovalbumin and BSA prior to washout. Histologic assessments at multiple timepoints included: 1) the presence of ILM-associated protein immunoreactivity measured as a linear distance over the explant surface and 2) a qualitative masked grading scale characterizing regularity and gaps in immunoreactivity (Supp Fig 3). In control BSS-treated retinal explants, laminin and collagen IV were present as a continuous band at the ILM that persisted unchanged through 11 days of culture. Both proteins were initially expressed within the retinal vasculature, though this diminished with time in culture (Fig 3A-C,E, Supp Fig 2A-C). Papain (10-45U/mL) and pronase E at the highest dose tested (3U/mL) eliminated laminin immunoreactivity at the ILM within hours, but laminin remained relatively preserved in retinal blood vessels and weakly in a few RGCL and INL cells suggesting spatial restriction of enzymatic activity (Fig 3A,E). In contrast, collagenase (20-30U/mL) and lower dose pronase E (0.6U/mL) caused subtle irregularity and discontinuity of laminin immunoreactivity at the ILM at day 0 (Fig 3A,E), and marked disruption by day 7 (Fig 3A,B,E, Supp Fig 2C).

**Figure 3.**
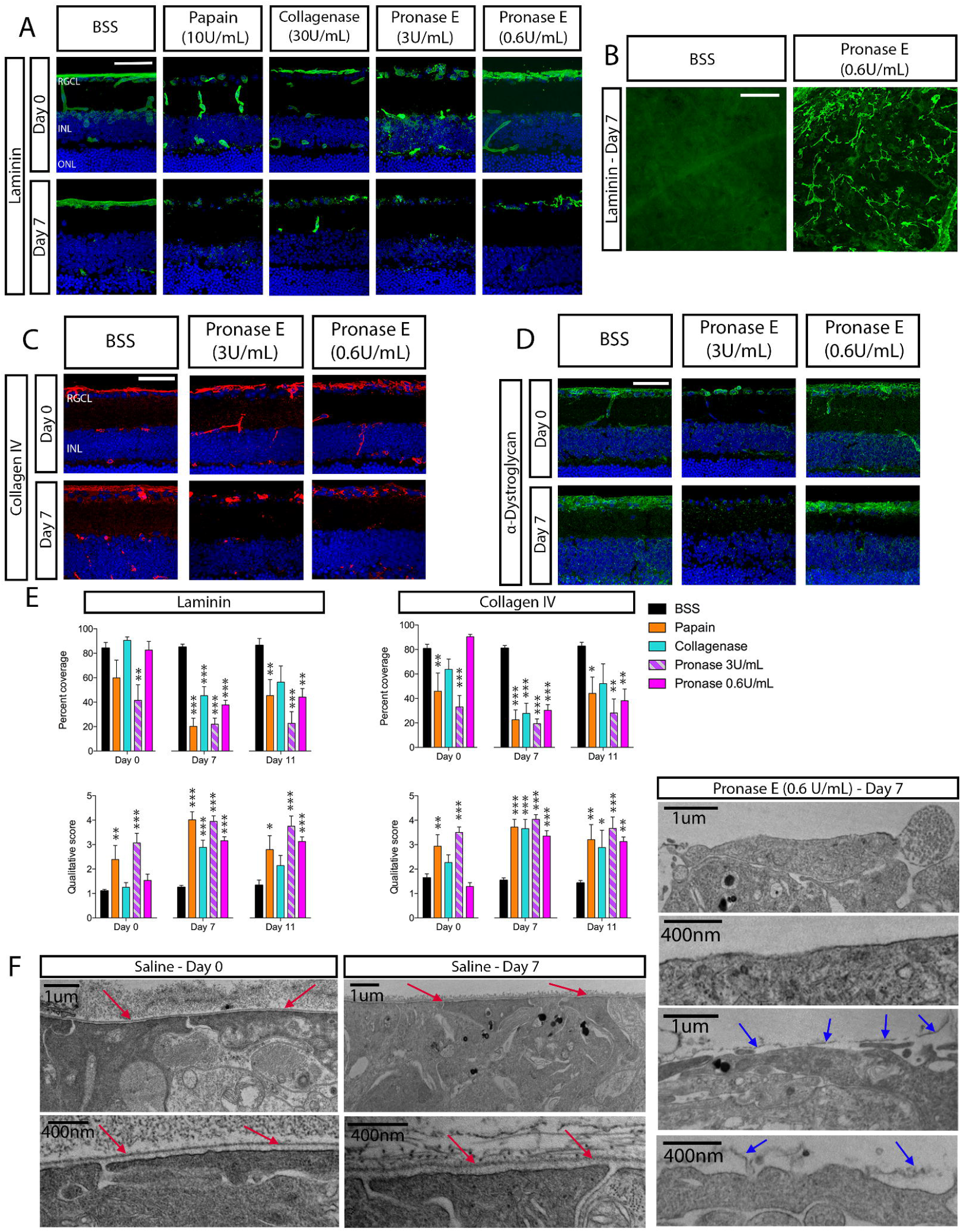
Effect of proteolytic enzymes on the internal limiting membrane. Adult mouse retinae were explanted and treated with proteolytic enzymes or basic salt saline (BSS, negative control) prior to inactivation and washout. Tissue was fixed within 1 hour, or after 7 or 11 days of organotypic culture. Laminin (A, B, green), collagen IV (C, red), and alpha-dystroglycan (D, green) were present at the ILM and expression was disrupted to varying degrees by enzyme treatment. Nuclei are counterstained with DAPI (blue). The immunofluorescence of laminin and collagen IV at the ILM were quantified using a qualitative grading scheme or by measuring the percent linear coverage of the retinal explant surface in cryosections, which showed ILM disruption by all enzymes tested (E). Transmission electron microscopy of the inner retinal surface reveals intact ILM over Müller glial endfeet in control retinal explants (red arrows), with overlying filaments of posterior vitreous cortex (F). The ILM was absent (top two micrographs) or fragmented (bottom two micrographs) following treatment with Pronase E (0.6 U/mL) (blue arrows), without alteration in the structure of the underlying retinal glia. Scalebars: 50μm (A-D) or as indicated (F). RGCL, retinal ganglion cell layer; INL inner nuclear layer; ONL, outer nuclear layer. Error bars: standard error of the mean (E). *p<0.05, **p<0.01, ***p<0.001 by one-way ANOVA with post-hoc Dunnett’s test for multiple comparisons versus the BSS control group.

Papain rapidly digested collagen IV at the ILM, whereas collagenase and both doses of pronase E caused irregularity and focal disruption of collagen IV staining at day 0 that were less extensive than those observed for laminin. After 7-11 days in culture, papain and pronase E (3U/mL) produced greater disruption in collagen IV reactivity at the ILM than collagenase or lower dose pronase E (Fig 3C,E, Supp Fig 2A,C).

Alpha-dystroglycan is a membrane-associated protein, localized to Müller cells at the ILM, that is important for cellular binding and signaling with ECM proteins including laminin.^38^ Alpha-dystroglycan immunoreactivity was strong at the ILM, but more granular in appearance than laminin or collagen IV in control retinal explants (Fig 3D, Supp Fig 2C). Papain, collagenase, and pronase E (3U/mL) were associated with early and persistent decreases in alpha-dystroglycan immunoreactivity at the ILM. However, following pronase E (0.6U/mL) treatment, alpha-dystroglycan immunoreactivity remained similar to control throughout the culture period (Fig 3D, Supp Fig 2B,C).

ILM degradation by proteolytic enzymes was confirmed by transmission electron microscopy. In BSS-treated retina, the ILM was a continuous electron dense linear membrane external to Müller cell footplates. Fibrillar deposits from adherent posterior vitreous cortex overlaid the ILM (Fig 3F). Papain, collagenase, and pronase (3U/mL) resulted in complete ILM loss, and also considerable degenerative changes to inner retinal Müller cells, astrocytes, and RGC axons (Supp Fig 3A-C). In contrast, pronase E (0.6U/mL) produced areas of near complete ILM loss juxtaposed to areas with focal ILM breaks, without significant changes to the underlying retinal ultrastructure (Fig 3F).

Retinal gliosis impairs intraretinal migration of transplanted MSCs.^44^ We sought to determine whether proteolytic ILM disruption also affected retinal viability or glial reactivity. Papain caused suppression of retinal gliosis and marked disruption of the laminar retinal architecture by 7 days in culture (Fig 3,4, Supp Fig 3). Collagenase and pronase E were associated with greater retinal laminar architecture preservation, though glial intermediate filament expression was suppressed after treatment with collagenase or pronase E (3U/mL) (Fig 4). The lower dose of pronase E (0.6U/mL), however, caused negligible change to the histological appearance of the retina and was associated with preservation of GFAP, vimentin, and nestin expression in astrocytes and Müller glia (Fig 4). Importantly, since transplanted RGCs would need to synapse with bipolar and amacrine cells, we documented that pronase E (0.6U/mL) resulted in negligible change to the ultrastructure of the IPL and INL at day 7 of culture (Supp Fig 4A-D).

**Figure 4.**
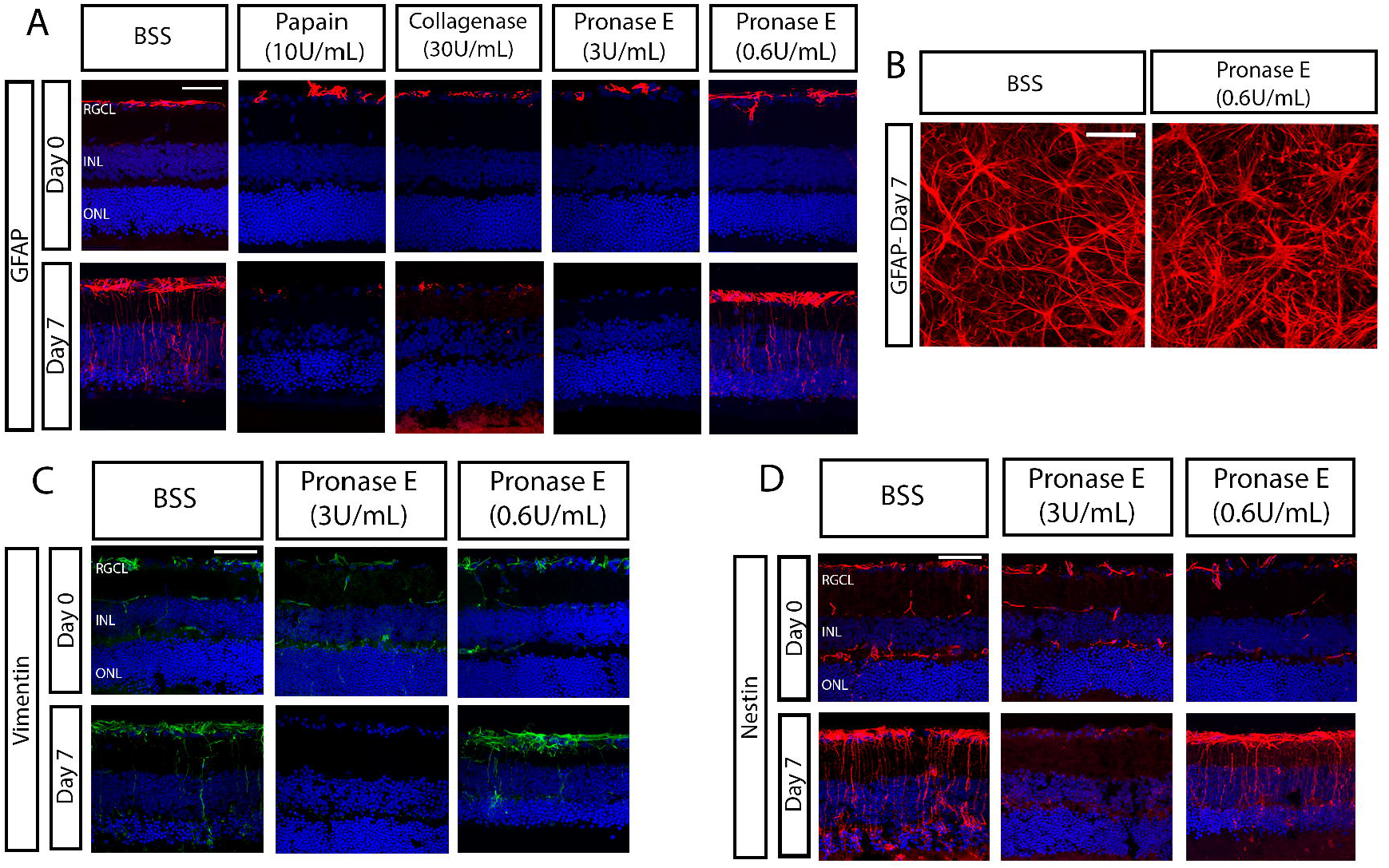
Effect of proteolytic enzymes on glial reactivity. Adult mouse retinae were explanted and treated with proteolytic enzymes or basic salt saline (BSS, negative control) prior to inactivation and washout. Tissue was fixed within 1 hour, or after 7 or 11 days of organotypic culture. Glial fibrillary acid protein (GFAP, A, B, red), vimentin (C, green), and nestin (D, red) were upregulated in astrocytes and Müller glia in BSS-treated explants as a result of organotypic culture. Nuclei are counterstained with DAPI (blue). Treatment with papain, collagenase, and Pronase E (3U/mL) resulted in suppression of reactive gliosis, whereas treatment with Pronase E (0.6U/mL) was associated with preservation of reactive gliosis. RGCL, retinal ganglion cell layer; INL, inner nuclear layer; ONL, outer nuclear layer. Scalebars: 50μm.

In sum, all enzymes tested caused marked and persistent ILM disruption. Papain caused unacceptable neurotoxicity at the concentrations tested. Collagenase and pronase E (3U/mL) caused less overt structural retinal degradation, but suppressed reactive gliosis. Pronase E (0.6U/mL) effectively digested the ILM without inducing detectable changes to the viability or physiology of the retina.

### Transplanted hES-RGC survival and topology following ILM disruption

In order to evaluate the effects of proteolytic ECM digestion on transplanted hES-RGCs, we applied and then inactivated and washed out enzymes ≥24h prior to transplantation (Fig 5A). Collagenase and pronase E (3U/mL) were tested because of their relatively mild effect on retinal explant architecture. However, they also negatively affected hES-RGC survival and were associated with high variability in topographic parameters, so further evaluation of these enzymes was not undertaken (Supp Fig 4A, Supplemental Results).

**Figure 5.**
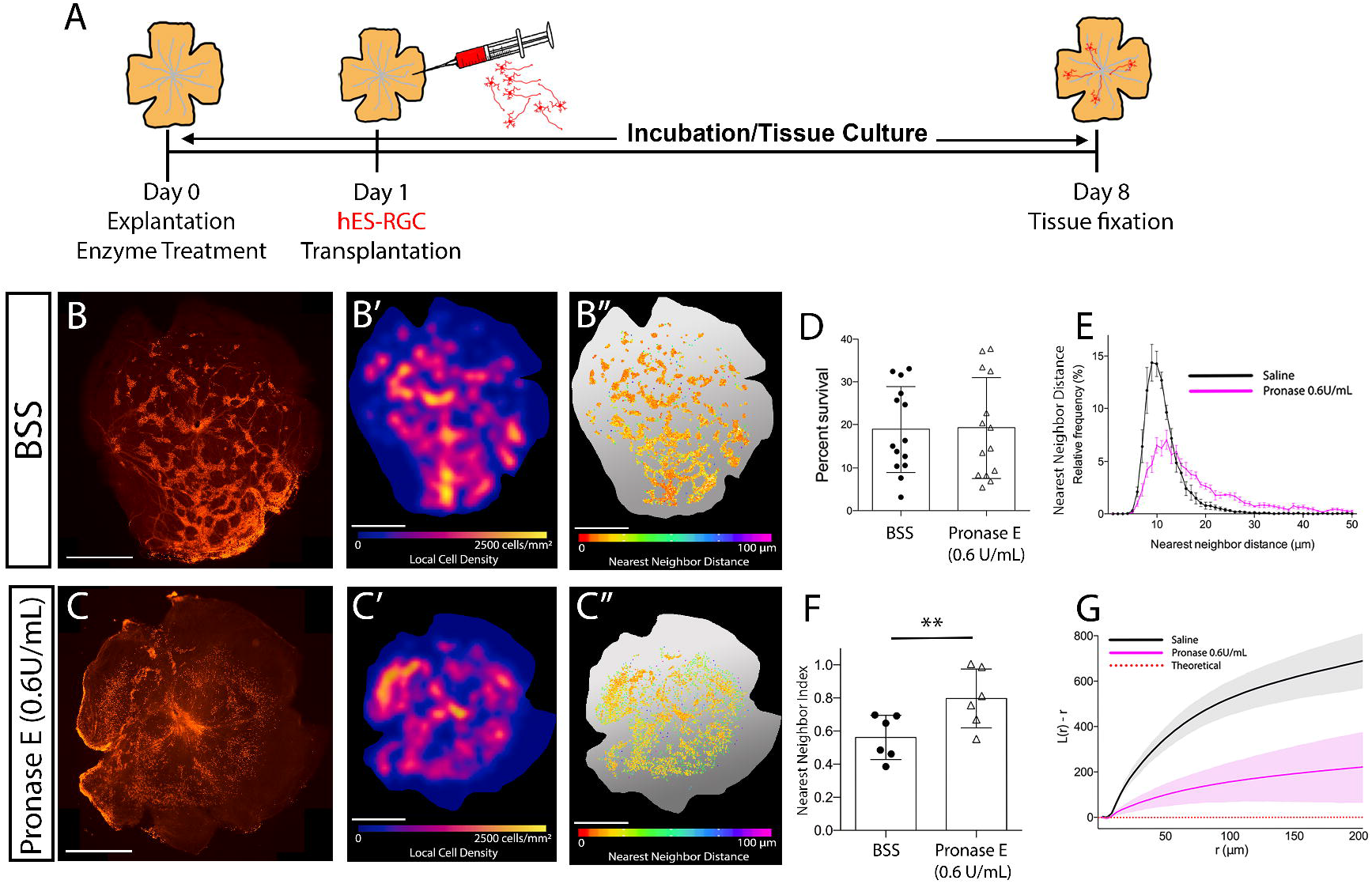
Topographic spacing of human embryonic stem cell derived retinal ganglion cells (hES-RGCs) transplanted onto retinal explants with or without proteolytic enzyme pre-treatment. The experimental paradigm is shown (A). Adult mouse retinae were explanted and treated with proteolytic enzyme or basic salt saline (BSS, negative control) prior to inactivation and washout. The following day, hES-RGCs were transplanted and co-cultured for 1 week prior to analysis. Epifluorescent micrographs show the tdTomato+ hES-RGC morphology (B, C). Cell density heatmaps (B’, C’) and nearest neighbor distance maps (B’’, C’’) demonstrate greater cell dispersal with enzyme pre-treatment. hES-RGC survival was similar in both groups (D). Nearest neighbor distance (E), nearest neighbor index (F), and Ripley’s L function (G) all demonstrated significantly less clustering following proteolytic enzyme treatment. Scalebars: 1.25mm. Error bars: standard deviation (D, F); standard error of the mean (E); 95% confidence interval (G). *p≤0.05, **p≤0.01, ***P≤0.001.

Consistent with its lack of apparent toxicity and in contrast to the other enzyme treatments tested, pronase E (0.6U/mL) resulted in no change in hES-RGC survival following transplantation compared to BSS (19.2±11.7% vs 18.9±10.0%, respectively, p=0.9, Fig 5D). Pronase E (0.6U/mL) was associated with dispersed rather than clustered hES-RGC spatial survival patterns (Fig 5B,C). Average hES-RGC density on pronase E-treated retinal explants was 210.2±147.1 cell/mm^2^ compared to 266.4±196.3 cell/mm^2^ for BSS (p=0.4). The average hES-RGC NN distance of pronase E-treated retinal explants was 24.2±7.2 μm vs 11.7±1.6 μm on control retinal explants (Fig 5E, p<0.01). The NNI was significantly higher in pronase E-treated explants compared to controls, indicating reduced clustering (Fig 5F). L(r)-r demonstrated a significantly attenuated rise over CSR for hES-RGCs on pronase E-treated retinal explants vs control, indicating reduced spatial clustering with enzymatic ECM disruption (Fig 5G). Nerve fiber bundling from hES-RGCs was not identified in pronase E (0.6UmL)-treated explants, unlike in controls (14.9 neurite bundles/100 hES-RGCs, Fig 5B,C).

In sum, compared to BSS-treatment, Pronase E (0.6U/mL) resulted in greater spatial dispersion of hES-RGCs with no clustering or neurite bundling, and without affecting overall survival, unlike collagenase or pronase E (3U/mL), which reduced hES-RGC survival.

### Transplanted hES-RGC neurite structural engraftment following ILM disruption

ECM digestion resulted in a dramatic increase in hES-RGC neurite localization into the retina after 7 days (Fig 6, Supp Fig 8). RGC somas, however, generally remained superficial to the retina or localized to the RNFL or RGCL without migrating deeper. Although collagenase and pronase E (3U/mL) did increase neurite penetration into the retina, the results were variable and of borderline statistical significance (Supp Fig 8, Supplemental Results).

**Figure 6.**
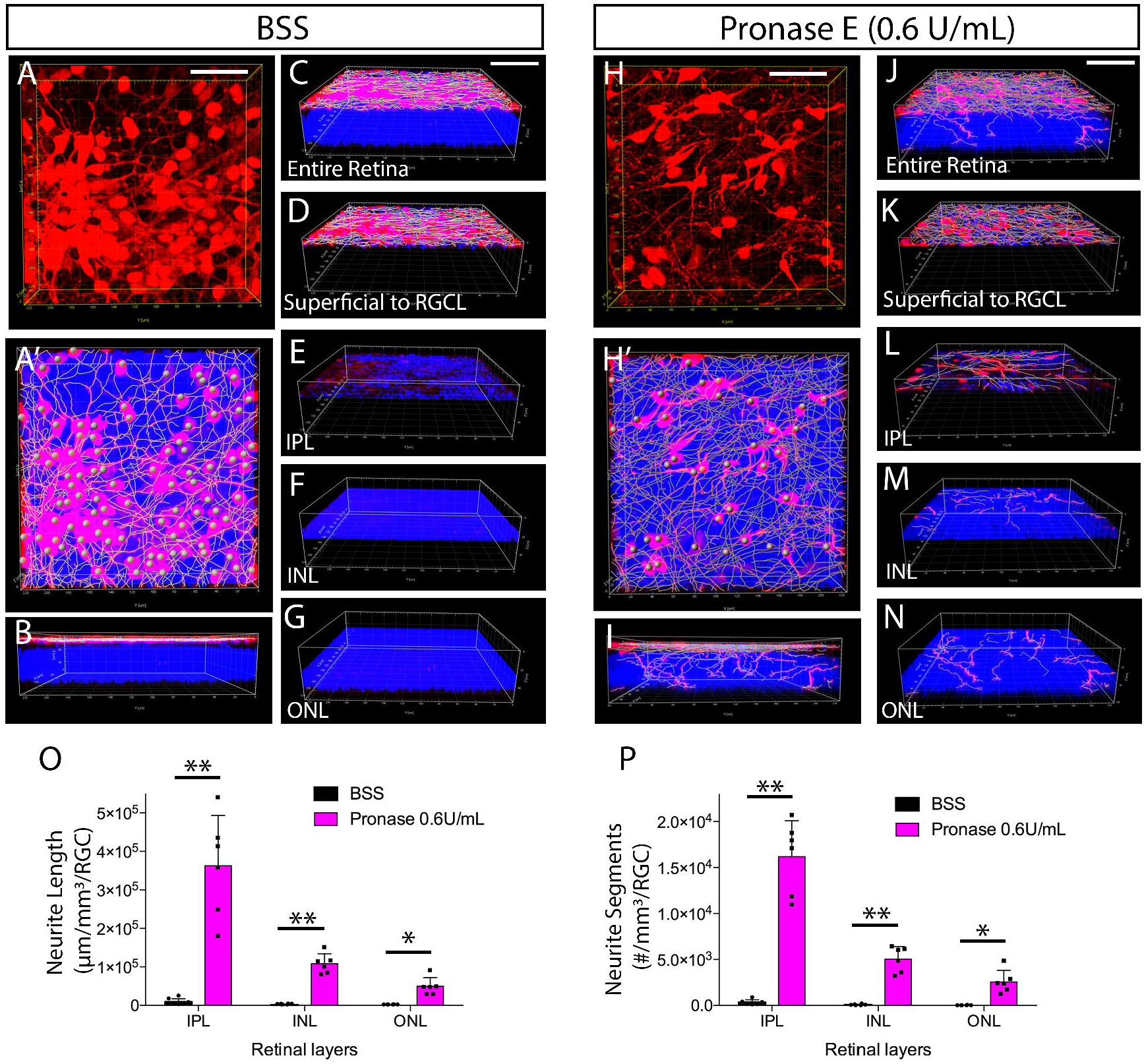
Retinal neurite ingrowth from human embryonic stem cell derived retinal ganglion cells (hES-RGCs) transplanted onto retinal explants with or without proteolytic enzyme pre-treatment. In saline (BSS, negative control) treated retinal explants, transplanted hES-RGCs (red) remained superficial to the neural retina, forming a distinct layer on top of the RGCL (A-G). Following pre-treatment with Pronase E (0.6 U/mL), hES-RGC neurites extended into the neural retina (H-N). Three dimensional reconstructions are shown (A-C, H-J) and the reconstructions were segmented according to retinal layer (D-G, K-N) to quantify hES-RGC neurite ingrowth on a spatially localized volumetric basis (O,P). RGCL, retinal ganglion cell layer; IPL, inner plexiform layer; INL, inner nuclear layer; ONL, outer nuclear layer. Scalebars: 50μm. Error bars: standard deviation (O, P). *p≤0.05, **p≤0.01 by unpaired t-test.

Interestingly, pre-treatment with pronase E (0.6U/mL) led to greater increase in hES-RGC neurite ingrowth into the retina than collagenase or pronase E (3U/mL), resulting in a more than 40-fold increase in the length and number of neurite segments in the IPL as compared to control explants (Fig 6, Videos 3,4). hES-RGC neurites could be found ectopically in the INL and ONL, though to a lesser extent than neurites located in the IPL. In pronase E (0.6U/mL)-treated retinal explants, there were 3.2-fold more hES-RGC neurites in the IPL than the INL and 6.3-fold more than in the ONL. Total neurite length was 3.2-fold greater in the IPL than the ONL, and 7.3-fold greater than in the ONL. We did not identify transplanted RGCs with dendrites that conformed to the morphology of any traditional RGC subtype.

To assess the possibility that material transfer of tdTomato RNA or protein to endogenous retinal neurons could have led to confusion about the source of the observed tdTomato^+^ neurites,^63–66^ we transplanted hES-RGCs onto retinal explants from ubiquitously-GFP expressing transgenic mice. We examined 337 hES-RGC somas superficial to the retina and 24 hES-RGC somas that had migrated into the retinal parenchyma, and 40 hES-RGC neurites within the retinal parenchyma using orthogonal confocal projections and fluorescence intensity histograms. We did not identify any presumed hES-RGCs with a neuronal morphology that co-expressed tdTomato and GFP (Fig 7E). Moreover, an antibody that specifically recognizes human nuclei labeled all tdTomato^+^ hES-RGC somas (Fig 7D), suggesting the tdTomato^+^ neurites visualized within the recipient retina arose from transplanted hES-RGCs.

**Figure 7.**
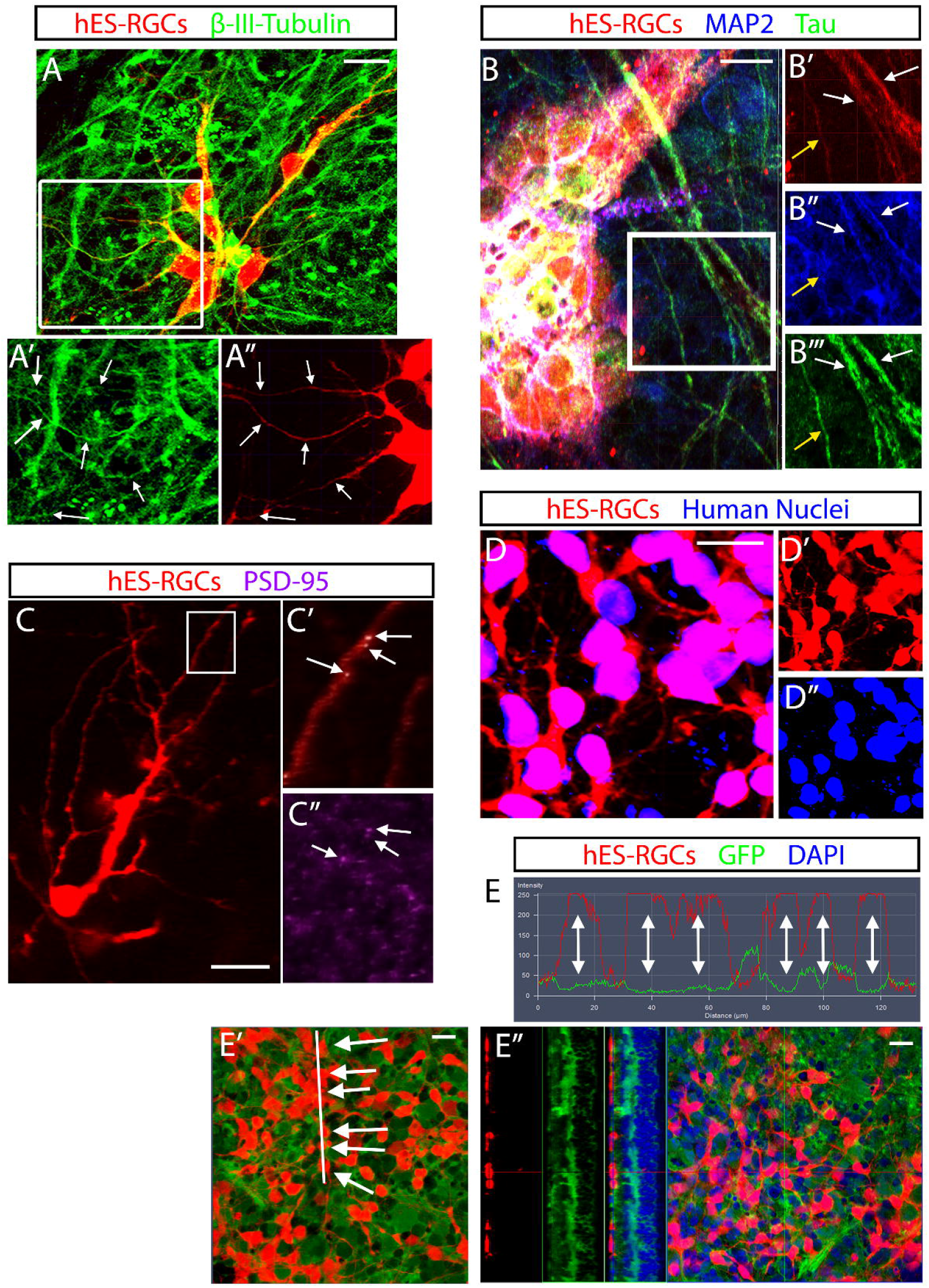
Characterization of structurally integrated hES-RGC neurites. One week after hES-RGCs (red) were transplanted onto retinal explants, immunofluorescence was used to characterize the transplanted neurons and their neurite processes. Transplanted hES-RGCs universally expressed β-III-tubulin (green) including in their neurites (A, arrows point to neurites co-expressing tdTomato and β-III-tubulin). Of note, β-III-tubulin is also expressed by the surviving endogenous RGCs which are tdTomato^−^. Neurites from hES-RGCs almost uniformly expressed Tau (green), but a subset co-expressed MAP2 (blue, B). White arrows in B indicate Tau^+^MAP2^+^ neurites, and the yellow arrow highights at Tau^+^MAP2^−^ neurite. Postsynaptic density-95 (PSD-95, purple, C’’) puncta could be found co-localizing with transplanted hES-RGC neurites within the inner plexiform layer (C). Colocalization confirmed by a thresholding algorithm using Imaris is indicated by white puncta (arrows, C’). hES-RGCs expressing tdTomato were uniformly labeled by antibody recognizing human nuclear antigen (D). hES-RGCs transplanted onto retinal explants isolated from transgenic mice ubiquitously expressing GFP were uniformly GFP-negative at the cell soma and within their neurite processes (E), as demonstrated orthogonal slices (E’’) and by immunofluorescence intensity histograms for tdTomato (red) and GFP (green) that demonstrate mutual exclusivity (E). Single-headed arrows (E’) point to cells tested for co-localization in the histogram shown, and double-headed arrows show the immunofluorescence profiles of those cells expressing tdTomato but not GFP (E). Scalebars: 20μm.

### Characterization of hES-RGC neurites within the neuroretinal parenchyma

We evaluated subcellular localization of canonical axonal or dendritic proteins in transplanted hES-RGCs. hES-RGCs universally expressed β-III-tubulin in the cell bodies and processes, regardless of location (Fig 7A). We next evaluated localization of Tau, which is expressed in mature axons, and MAP2, which is expressed in mature dendrites (Fig 7B). Of note, immature developing neurons segregate MAP2 and Tau only after specification of the axon.^67^ Therefore, developing neurons may co-localize these proteins within immature neurites. Within 296 individual neurite processes, we identified a differential expression pattern of MAP2 and Tau that correlated with neurite localization within the host retina. On the retinal surface, Tau and MAP2 were co-expressed in 92.7% of hES-RGC neurites, whereas 6.9% of neurites expressed Tau only. Within the retinal parenchyma, however, 73.5% of neurites co-expressed Tau and MAP2 (p=0.001 vs surface), 20.4% expressed Tau only (p=0.005 vs surface), and 4.1% expressed MAP2 only (p=0.069 vs surface). This observation might suggest that structural localization within the host retina promotes the maturation of hES-RGCs or that localization into the neuroretina follows neurite specification. However, Tau^+^ axons localizing deep to the RNFL would be ectopic.

To determine the propensity of transplanted RGC neurites to synapse with recipient retinal neurons, we used immunofluorescence to visualize synaptic proteins colocalizing with hES-RGC neurites. The postsynaptic protein PSD-95 was present in puncta that occasionally colocalized with hES-RGC somas and neurites (Fig 7C). We also identified examples of neurites in the deep retinal layers that expressed PSD-95 in a punctate pattern, consistent with what one would expect from a mature integrated RGC dendrite.

## Discussion

While human stem cell-derived RGCs extend neurites following transplantation onto retinal tissue, those neurites do not spontaneously localize within the retinal parenchyma. Our data suggest that the ILM may be a major barrier to neurite engraftment, as mechanical disruption or enzymatic degradation of ILM proteins were associated with marked increases in retinal neurite ingrowth. Spatial localization of RGC dendrites within the IPL is necessary for afferent synaptogenesis, and developing methods to permit donor RGC neurites to bypass the ILM is critical to the future of RGC replacement. Given the low efficiency of transplanted RGC engraftment documented in the most encouraging work to date,^29^ our data provide a clear avenue towards improving RGC transplantation outcomes in pre-clinical models.

### Transplanted RGC neurite engraftment

In untreated retinal explants, we identified virtually no interaction between transplanted RGCs and underlying retinas except in locations with physical disruption to the retina and ILM. It is noteworthy that the organotypic retinal explant culture model likely biases towards greater engraftment than intravitreal injection in vivo, given that transplanted neurons are maintained in direct opposition to the ILM rather than being suspended within the vitreous cavity. Therefore, the exclusion of hES-RGCs from integration under control conditions supports the validity of this system in modeling barriers to intravitreal transplantation.

The ILM limits neuroretinal transduction by intravitreally-administered AAV vectors. Experimental enzymatic digestion,^32^ surgical ILM peeling,^30,68^ and sub-ILM injection^69^ circumvent this barrier, though the latter two are likely not feasible in rodent eyes. Here, we demonstrate that enzymatic digestion of ECM proteins within the ILM enhances retinal neurite ingrowth of transplanted RGCs. From a translational perspective, addressing the ILM barrier will be necessary to achieve functional RGC replacement. Given that ILM thickness increases substantially with age and in the presence of common diseases like diabetes,^70^ patients suffering from age-related optic neuropathies such as glaucoma and ischemic optic neuropathy may have a greater impediment to engraftment. Enzyme administration may not be necessary in humans, since surgical ILM peeling is a well-established and safe maneuver for the treatment of macular hole.^71^ Indeed, it is also possible that intravitreal proteolytic enzymes would be toxic at the concentrations needed to digest the ILM clinically. For example, we found that intravitreal papain, pronase E, and collagenase at the concentrations used here digested the retinal vasculature and induced intraocular hemorrhage in living mice (data not shown). Though additional experiments are necessary to evaluate the role of the ILM in transplanted hES-RGC engraftment *in vivo,* such experiments may require development of alternative methodologies for ILM disruption.

We have previously assessed barriers to retinal integration of bone marrow derived MSCs and noted that whereas ILM digestion with collagenase did not permit transplant integration, suppression of reactive gliosis with alpha-aminoadipic acid did.^44^ In that case, MSCs entered the retina in spite of an intact ILM, suggesting that this structure is not a generalized physical barrier. It is therefore notable that for hES-RGCs, ILM digestion that specifically preserves glial reactivity dramatically improves structural cell integration. We speculate that differential effects of the ILM and retinal gliosis on engraftment of these two cell types is modulated by differential expression of cell surface receptors that mediate interactions with the ECM and glia. Identification of surface receptors mediating hES-RGC interactions with the ILM could therefore inform methods of permitting trans-ILM retinal integration without the need for disrupting the ILM directly, and is a subject of ongoing investigation. Indeed, given the importance of RGC-ILM interactions for retinal patterning during development, total disruption of the ILM may in fact be counterproductive. For instance, signaling between developing RGCs and laminin within the ILM is involved in polarity decisions and axon localization to the basal retina, though polarization does persist in the absence of this signaling at a delayed pace.^72^

We noted that hES-RGC neurites entering the retinal parenchyma did not exclusively target the IPL. The developmental factors that control RGC dendrite laminar patterning to and within the IPL include both molecular cues and activity-dependent refinement.^73^ Spontaneous electrophysiological activity in organotypic retinal explants is modest, decreases with time, and may have limited any potential IPL-directed dendrite localization reinforced by neuronal activity.^74^ During development, RGC dendrites target pre-patterned IPL afferents.^75^ Sublamination within the IPL is guided by the expression of specific cell surface receptors and their binding to localized lamina-specific ligands, including integrins, cadherins, plexins, and others play critical roles in dendritic outgrowth and guidance.^35,76^”^83^ It is conceivable that controlling expression of relevant surface receptors may aid in localizing hES-RGC dendrites to specific locations where afferent synaptogenesis may occur. It remains unclear the extent to which ligand expression is maintained within the mature IPL, but the identification of appropriate dendritic stratification by transplanted primary RGCs suggests that at least some of the necessary signals remain present.^29^

### Transplanted RGC Survival

Consistent with prior reports documenting limited survival and integration of transplanted RGCs,^26,29^ we observed hES-RGC survival rates of 10-30% at one week. Survival was not affected by retinal pre-treatment with pronase E (0.6U/mL), though it was further reduced by pronase E (3U/mL) or collagenase, which also impaired retinal glial activity. Our finding that transplanting fewer (1.5×10^4^) hES-RGCs was associated with inferior survival compared to transplantation of 2.5×10^4^ or 5×10^4^ cells is in contrast to the report by Venugopalan et al.^29^ showing that transplanting 4×10^4^ RGCs resulted in 3-fold greater survival than 6×10^4^ cells. Differences in the source of transplanted RGCs, the recipient species and model system, or the experimental time period following transplantation may explain these findings. Alternatively, the association between transplanted RGC number and survival rate may be represented by an inverted U-shaped curve, where too few or too many transplanted cells are suboptimal. Regardless, improving RGC survival following transplantation is a key goal for ongoing research.

### Transplanted hES-RGC topographical localization

We observed a stark difference in spatial clustering of transplanted hES-RGC somas with and without proteolytic enzyme pre-treatment of the recipient retina. Cell clumping on retinal explants, a phenomenon that was not observed in cell culture, was attenuated by ECM digestion. This might suggest that interactions between the hES-RGCs and the ILM promote cell clustering. It is interesting, however, that transgenic disruption of neuronal interactions with the ILM through dystroglycan,^38^ integrin-β1, or Cas adaptor proteins^35^ results in ectopic clustering of RGCs and amacrine cells on the basal retinal surface. In order to quantify our observation in a statistically robust manner, we employed a number of spatial analytic tools including NN distance distributions (which reflect hyperlocal cell relationships between only adjacent cells), DRPs (which provide insight into densities over larger distances), and Ripley functions (cumulative functions that facilitate normalization to CSR conditions and therefore comparisons between experimental groups). The relative strengths and weaknesses of these tools for characterizing retinal neuron mosaicism have been elegantly reviewed recently.^84^

Unlike the clear necessity for hES-RGC dendrites to be proximally localized to bipolar and amacrine processes for synaptogenesis, the significance of hES-RGC soma clustering on intact ILM is unclear. The NN distance of hES-RGCs within clusters was only marginally lower than that of packed endogenous RGCs, and high-density coverage of the retina may be necessary to obtain high resolution retinotopic physiology. But lateral spreading of transplanted hES-RGCs will be necessary to achieve widespread coverage. Whether somal clumping and neurite ingrowth are directly related will need to be determined by future work.

### Material Transfer

A critical aspect of this work was the exclusion of “material transfer” as a possible explanation for the presence of tdTomato^+^ neurites within the host retina. Early photoreceptor transplantation experiments were interpreted as demonstrating a high degree of donor cell integration. However, subsequent experiments demonstrated that labeled donor photoreceptors transplanted into the subretinal space transfer either label RNA or protein to host cells through still-unclear mechanisms, and that most labeled cells in the ONL are actually host-derived and secondarily acquire label through material transfer.^63–66^ Whereas RGC participation in material transfer has not been reported, it is prudent for all neuronal transplantation work to include control experiments to assess for this phenomenon. In this work, tdTomato^+^ cells with neuronal morphology transplanted into pan-GFP mice did not express GFP, indicating that they were not host-derived. Further, they uniformly expressed human nuclear antigen, consistent with their donor origin.

### Limitations

There are several limitations of the organotypic retinal explant system employed in this work. There is a temporal limit to the viability of the host tissue of about 10-14 days in cultures, which restrict experimental duration of less than might be required for functional synaptogenesis. The expression of important chemotactic or inhibitory factors may change in retinal explants as compared to in vivo retina, as this has not been specifically evaluated. There is no circulation or immune system, so immunologic rejection could not be modeled. The challenge of effective ILM disruption in rodents *in vivo* without disruption of other retinal processes has been discussed. As such, future work conducted *in vivo* will need to overcome numerous additional obstacles to the integration of transplanted RGCs into the retinal neurocircuitry. However, the identification of the ILM as a primary barrier will be critical to those future experiments and the data provided here suggest an approach to increase the efficiency of transplanted RGC engraftment. This work also does not address axon outgrowth, pathfinding, or efferent synaptogenesis. Such studies await efficient engraftment of transplanted RGCs in an *in vivo* model.

### Conclusions

We have characterized the spontaneous morphologic behavior of hES-RGCs transplanted onto mammalian neurosensory retina. Transplanted cells demonstrate clustering of cell somas, bundling of nerve fibers, and exclusion of neurites from the retinal parenchyma. Proteolytic digestion of ECM proteins within the ILM is associated with reduced cell body clustering, a lack of fiber bundling, and a profound increase in neurite ingrowth into the retina. It is likely that modifying the interactions between transplanted RGCs and the ILM will be necessary to facilitate efficient functional engraftment for RGC replacement and optic nerve regeneration.

## Supporting information

Graphical Abstract

## Author Contributions

KYZ: Collection/assembly of data, data analysis & interpretation, manuscript writing, final manuscript approval

CT: Collection/assembly of data, data analysis & interpretation, manuscript writing, final manuscript approval

JLM: Provision of study material, manuscript writing, final approval of manuscript

SQ: Collection and assembly of data, manuscript writing, final approval of manuscript

LW: Data analysis and interpretation, final approval of manuscript

HAQ: Financial support; provision of study material; data analysis & interpretation; final approval of manuscript

DLZ: Financial support; provision of study material, data analysis & interpretation, final approval of manuscript.

TVJ: Conception and design; financial support, administrative support, provision of study material, collection and assembly of data, data analysis and interpretation, manuscript writing, final approval of manuscript

## Acknowledgements

This work was funded by: the National Eye Institute (K12EY015025 [TJ]; R01EY002120 [HQ]; P30EY001765 [DZ]); The ARVO David L Epstein Award [HQ and TJ]; Research to Prevent Blindness (Career Development Award [TJ] and Unrestricted Grant Funding [Wilmer Eye Institute]); The American Glaucoma Society (Mentoring for the Advancement of Physician Scientists Grant [TJ]); The Johns Hopkins Physician Scientist Training Program (Pilot Grant [TJ]); and generous gifts from the Guerrieri Family Foundation, the Gilbert Family Foundation, and the Marion & Robert Rosenthal Family Foundation. The authors are grateful to Dr. Alex Kolodkin (Dept of Neuroscience, Johns Hopkins University) for critical evaluation of the manuscript.

## Data Availability Statement

Primary data are available upon request to the corresponding author.

## Supplemental Methods

### Human stem cell derived RGCs

For differentiation to an RGC fate, hES cells were enzymatically dissociated to single cells using Accutase (Millipore-Sigma, Burlington, MA) and re-plated on Matrigel in mTeSR-1 containing blebbistatin (Millipore-Sigma). On the following day, designated day zero, cells were switched to 5% CO_2_/20% O_2_ and the culture media was replaced with 1:1 DMEM/F12 and Neurobasal plus 1% N2 Supplement, 2% B27, 1% GlutaMAX, and 1% antibiotic/antimycotic (all from ThermoFisher Scientific, Carlsbad, CA), which was fully exchanged at least every other day. To increase differentiation efficiency, small molecules were added to the media: 1μM Dorsomorphin and 2.5μM IDE2 (both from R&D Systems, Minneapolis, MN) from day 1-6, 10mM nicotinamide (Millipore-Sigma) from days 1-10, 25μM forskolin (Cell Signaling Technology, Danvers, MA) from days 1-30, and 10μM DAPT (Cell Signaling Technology) from days 18-30. On day 35-40, cells were dissociated using Accumax (Millipore-Sigma) and purified using the anti-CD90.2/Thy1.2 MACS system (Miltenyi Biotec, Auburn, CA), following the manufacturer’s protocol.

After purification, some hES-RGCs were frozen at 1×10^7^ cells/mL in CryoStor CS10 (StemCell Technologies) by centrifuging cells at 150g for 6min, suspending the pellet in chilled CS10 in a cryovial and freezing in controlled-rate freezing container at −80°C overnight before transfer to liquid nitrogen (−130°C) for storage. hES-RGCs were thawed by placing cryovials in a 37°C bath then diluting 10:1 with retinal explant culture media. The cell suspension was equilibrated for at least 3h in a tissue culture incubator a loose lid facilitating gas exchange, then centrifuged at 300g for 10min, and the pellet was resuspended in retinal explant culture media.

### Organotypic retinal explants

Retinas were isolated following mouse euthanasia by overdose of intraperitoneal ketamine and xylazine. When retinal explants were cultured without hES-RGC transplantation, some retinas were sectioned into radial quarters at explantation; otherwise retinas were explanted whole in Maltese cross configuration with radial relaxing incisions. In some cases, following fixation, explants were cut in half prior to histology. Retinal explants were cultured at an air-fluid interface overlying culture media composed of Neurobasal-A, B27 supplement (2%), N2 supplement (1%), L-glutamine (0.8mM), penicillin (100U/mL), and streptomycin (100μg/mL, all from ThermoFisher Scientific). Cultures were maintained in dark, humidified conditions at 37°C in 5% CO_2_. Half of the media was exchanged every other day.

### Proteolytic enzymes

Papain (Worthington Biochemical Corp, Lakewood NJ) was reconstituted in balanced salt saline (BSS) with 1.1mM EDTA, 0.067mM mercaptoethanol and 5.5mL cysteine-HCl. Pronase E from *Streptomyces griseus* (Millipore-Sigma) and collagenase type I-A from *Clostridium histolyticum* (Millipore-Sigma) were reconstituted in BSS. Proteolytic enzymes were prepared fresh from powder substrate, 0.22μm syringe filtered, and incubated at 37°C for 1h before each use.

### Immunofluorescence

Retinal explants were fixed by immersion in cold 4% paraformaldehyde for 12h. To obtain cryosections, tissue was washed in PBS and then incubated in 25% sucrose in PBS for 24h prior to freezing in TissueTek® OCT compound (Sakura Finetek USA, Inc., Torrance, CA) on dry ice. Cryosections were cut to 16μm thickness. Immunofluorescence was performed on cryosections or on flatmount retinal explants. Tissue was washed twice in PBS and then simultaneously blocked and permeabilized in PBS with 5% normal goat serum (NGS) for cryosections or 10% NGS (flatmounts) and 0.1% Triton at room temperature (1 hour for sections, 3 hours for flatmounts). Primary antibody was diluted in blocking solution (Supp Table 1) and tissue was immersed at 4°C (12h for sections, 48h for flatmounts on a shaker). Secondary antibody was diluted in blocking solution (Supplemental Table 1) and tissue was immersed at room temperature (3h for sections, 24h for flatmounts on a shaker). Nuclei were counterstained with DAPI (Millipore-Sigma, 0.1μg/mL). Tissue was cover-slipped under immunofluorescence mounting media (Dako North America, Carpinteria, CA).

Some primary antibodies were optimized by special processing. Vimentin immunofluorescence included treatment with pepsin reagent antigen retriever (Millipore Sigma) for 5min at 37°C. followed by three PBS washes, permeabilization in PBS with 0.25% Triton for 30min at room temperature, and blocking with 5% NGS and 0.1% BSA for 90min. Alpha-dystroglycan immunofluorescence included permeabilization with 0.25% Triton in PBS for 10min, followed by blocking with 2% NGS in PBS with 0.1% BSA for 30min. Collagen IV immunofluorescence included antigen retrieval with buffer containing 10mM sodium citrate (pH 6.0) and 0.05% Tween-20 at 95°C for 3min. Slides were cooled to room temperature, washed and permeabilized in PBS with 0.25% Triton, and blocked in 5% NGS and 0.1% BSA in PBS.

### Transmission electron microscopy

Retinal explants were immersion fixed in a 2% paraformaldehyde/2.5% glutaraldehyde in Cacodylate buffer. Tissue was post-fixed in 1% osmium tetroxide for 2h, dehydrated in ascending alcohol concentrations, and stained in 1% uranyl acetate in 100% ethanol for 1h. Tissues were embedded in epoxy resin mixture at 60°C for 48h. One micron sections were cut and stained with 1% toluidine blue. Ultra-thin sections (~68nm thick) were collected on copper grids, stained with uranyl acetate and lead citrate, and imaged with a Hitachi H7600 transmission electron microscope (TEM, Hitachi High Technologies, Clarksburg, MD).

### Image analyses

For RGC quantification and spatial density measurements, RGCs soma locations were delineated using the multi-point selection tool and retinal area was traced using the freehand selection tool. For topography analyses, a separate retinal edge was defined 150μm inside the true tissue edge, and RGCs outside of the topography analysis edge were excluded to avoid artifactual clustering of RGCs at the boundary of the transplantation space, since RGCs could not be located past the tissue edge. Topography of endogenous RGCs, immunohistochemically labeled with antibody against RBPMS, was assessed similarly.

Local cellular topography was assessed by NN distances and NN index. Each cell has a single value for NN distance whereas the NN index is calculated for each retinal explant. NN index equals the average NN distance observed divided by, the theoretical NN distance under conditions of complete spatial randomness (CSR), where 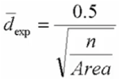. Here, *n* is the number of RGCs in an explant, and *Area* is the surface area of the retinal explant convex hull occupied by RGCs, which may be distinct from the total retinal area including or excluding the retinal edge. Density heat maps of RGC point patterns were generated using the density.ppp function from spatstat to compute a kernel smoothed intensity graph. NN distance heat maps were generated using the ggplot2 R package.

## Supplemental Results

### Transplanted hES-RGC survival and topology following ILM Disruption

hES-RGCs survival on retinal explants pretreated with collagenase 30U/mL and pronase E (3U/mL) was lower (5.8±2.4% and 10.2±4.0%, respectively) than on control retinas (15.6±5.4%, p<0.05). In general, enzyme pre-treatment of retinal explants was associated with reduced clustering of hES-RGC somas and greater spatial regularity following transplantation (Supp Fig 4A-C). The average NN distance in collagenase and pronase E (3U/mL) treated explants (31.2±13.6μm and 22.0±4.8μm, respectively) were significantly higher than the NN distance of control explants (15.2±1.7μm, p<0.05). However, hES-RGC survival was poor and topology was highly variable in collagenase and pronase E (3U/mL) treated explants; therefore we elected not to characterize the spatial properties in these cultures further.

### Transplanted hES-RGC neurite structural engraftment following ILM disruption

Compared with control retinal explants (Supp Fig 8A-D), pre-treatment with pronase E (3U/mL) resulted in an increase in hES-RGC neurite ingrowth into the IPL of almost 15-fold (Supp Fig 8E-H). Of note, hES-RGC neurites also extended to ectopic locations within the INL and ONL of enzyme-treated retinal explants, though to a lesser extent than the IPL (Supp Fig 8I-J). We did not identify transplanted RGCs that conformed to the morphology of any traditional RGC dendritic branching pattern. Collagenase pre-treatment also increased the number and length of hES-RGC neurites that penetrated the neural retina, though the effect was variable across explants and not statistically significant overall (Supp Fig 8I-J).

## Supplementary Figure Legends

**Supplementary Figure 1. Topographic spacing of cryopreserved and thawed hES-RGCs in culture.** Human embryonic stem cell (hES) derived retinal ganglion cells (RGCs) were cryopreserved, thawed, and then cultured on poly-L-ornithine and laminin-coated for 1 week. Epifluorescence microscopy revealed the morphology and spacing of tdTomato^+^ hES-RGCs (A). A heat map (A’) shows local cell density and the nearest neighbor distance map (A’’) shows the distance between each cell and his nearest neighbor, which is also plotted as a histogram (B). The density recovery profile (DRP, C) demonstrates the mean RGC density as a function of distance from each RGC in the sample. Scalebars: 1.25mm. Error bars: standard error of the mean (B, C).

**Supplementary Figure 2. Effects of proteolytic enzymes on the internal limiting membrane.** Cryosections of retinal explants cultured for 0, 7, or 11 days are shown. Proteolytic enzymes were applied in the indicated concentrations at day 0. Collagen IV (A, C, red), α-dystroglycan (B, C, green), and laminin (C, green) were disrupted at the ILM by various enzyme digestions. Nuclei are counterstained in blue with DAPI. RGCL, retinal ganglion cell layer; INL, inner nuclear layer. Scalebars: 50μm.

**Supplementary Figure 3. Qualitative grading system for internal limiting membrane disruption.** A qualitative system for characterizing the degree of retinal explant ILM disruption was developed, whereby a masked investigator scored micrographs of cryosections processed for immunofluorescence against laminin and collagen IV. Representative imagines of explants for each score are shown (A). Grade 1: Strong, continuous, linear immunofluorescence at the ILM throughout the explant; Grade 2: Linear immunofluorescence at the ILM with small disruptions present; Grade 3: Linear immunofluorescence at the ILM with large disruptions present; Grade 4: Markedly segmented, disjointed immunofluorescence at the ILM with few or no continuous linear segments; Grade 5: Little to no immunofluorescence at the ILM. A scatterplot comparing qualitative ILM immunofluorescence scores with average percent coverage scores for individual explants (average of at least three separate micrographs per explant) demonstrated a strong inverse correlation for both laminin and collagen IV (B).

**Supplementary Figure 4. Transmission electron microscopy of the internal limiting membrane of retinal explants with or without pre-treatment with Papain (10 U/mL), Pronase E (3 U/mL) or Collagenase (20 U/mL).** Retinal explants that had been treated with the indicated proteolytic enzymes at day 0 and then cultured for 7 days demonstrated total digestion of the internal limiting membrane, but there were also extensive degenerative changes evident to the retinal cells at the inner retinal surface. Scalebar: 1μm.

**Supplementary Figure 5. Effects of proteolytic enzymes on the retinal gliosis.** Cryosections of retinal explants cultured for 0, 7, or 11 days are shown. Proteolytic enzymes were applied in the indicated concentrations at day 0. Vimentin (A, C, green), Nestin (B, C, red), and glial fibrillary acid protein (GFAP, C, red) were upregulated during retinal explant culture with BSS or Pronase E (0.6 U/mL) treatment, but not with the other enzymes tested. Nuclei are counterstained in blue with DAPI. RGCL, retinal ganglion cell layer; INL, inner nuclear layer; ONL, outer nuclear layer. Scalebars: 50μm.

**Supplementary Figure 6. Transmission electron microscopy of the inner nuclear and plexiform layers of retinal explants with or without pre-treatment with Pronase E (0.6 U/mL).** Retinal explants that had been cultured for 7 days demonstrated preservation of inner nuclear layer morphology and neurite ultrastructure integrity within the inner plexiform layer when pretreated with either saline (BSS, negative control, A, C) or Pronase E (0.6 U/mL, B, D) at day 0.

**Supplementary Figure 7. Topographic appearance of hES-RGC transplanted onto retinal explants with or without pre-treatment with Pronase E (3 U/mL) or Collagenase (20 U/mL).** Adult mouse retinae were explanted and treated with proteolytic enzyme (B, C) or basic salt saline (BSS, negative control, A) prior to inactivation and washout. The following day, hES-RGCs were transplanted and co-cultured for 1 week prior to analysis. Epifluorescent micrographs show the tdTomato+ hES-RGC morphology. Scalebars: 1.25mm.

**Supplementary Figure 8. Structural integration of transplanted hES-RGC neurites into retinal explants with or without pre-treatment with Pronase E (3 U/mL) or Collagenase (20 U/mL).** In saline (BSS, negative control) treated retinal explants, transplanted hES-RGCs (red) remained superficial to the neural retina, forming a distinct layer on top of the RGCL (A-D). Following pre-treatment with Pronase E (3 U/mL), hES-RGC neurites extended into the neural retina (E-H). Three dimensional reconstructions are shown (A, E) and the reconstructions were segmented according to retinal layer (B-D, F-H) to quantify hES-RGC neurite ingrowth on a spatially localized volumetric basis (I, J). Maximum projection images of segmented retinal layers are shown (D, H). RGCL+, superficial surface transplant layer through the retinal ganglion cell layer; IPL, inner plexiform layer; INL, inner nuclear layer; ONL, outer nuclear layer. Scalebars: 50μm. *p≤0.05, **p≤0.01 by one-way ANOVA with post-hoc Dunnett’s test in comparison to the BSS group.

## Supplementary Videos

**Supplementary Video 1: 3D animation showing transplanted hES-RGCs at the center of a retinal explant.**

Transplanted hES-RGCs (red) form a monolayer on the surface of and external to the underlying organotypic retinal explant. Nuclei are labeled with DAPI (blue).

**Supplementary Video 2: 3D animation showing transplanted hES-RGCs at the edge of a retinal explant.**

Transplanted hES-RGCs (red) located near the edge of retinal explants or at radial relaxing incisions (bottom right at the beginning of the movie) enter the retinal tissue at the cut edge and send neurites that migrate laterally through the retina. Nuclei are labeled with DAPI (blue).

**Supplementary Video 3: 3D animation showing transplanted hES-RGCs on a BSS-treated retinal explant.**

Transplanted hES-RGCs (red) form a monolayer on the surface of and external to the underlying organotypic retinal explant after treatment with BSS (negative control). Nuclei are labeled with DAPI (blue). Cell bodies were manually tagged (balls) and neurites semi-manually traced (grey filaments).

**Supplementary Video 4: 3D animation showing transplanted hES-RGCs on a Pronase E-treated retinal explant.** Following pre-treatment of the recipient retinal explant with pronase E (0.6U/mL), transplanted hES-RGCs (red) send numerous neurite filaments into the underlying retinal parenchyma. The majority localized to the inner plexiform layer but off-target neurites projecting into the inner and outer nuclear layers are also seen. Nuclei are labeled with DAPI (blue). Cell bodies were manually tagged (balls) and neurites semi-manually traced (grey filaments).

