## Supplementary Table 1 for "Structural engraftment and topographic spacing of transplanted human stem cell-derived retinal ganglion cells"

| **Antigen Target** | **Host Species** | **Clonality** | **Clone** | **Dilution** | **Company** | **Catalog Number** |
| --- | --- | --- | --- | --- | --- | --- |
| Red fluorescent Protein (RFP, cross-reactive to tdTomato) | Rabbit | Polyclonal | NA | 1:200 | Rockland | 600-401-379 |
| RFP, cross-reactive to tdTomato | Mouse | Monoclonal IgG1 | RF5R | 1:500 | Thermo-Fisher | MA5-15257 |
| RFP, cross-reactive to tdTomato | Goat | Polyclonal | NA | 1:500 | Sicgen | AB8181-200 |
| Human Nuclear Antigen (HuNu) | Mouse | Monoclonal IgG1 | 235-1 | 1:500 | Millipore-Sigma | MAB1281 |
| Laminin | Rabbit | Polyclonal | NA | 1:1000 | Millipore-Sigma | L9393 |
| Collagen IV | Mouse | Monoclonal IgM | J3-2 | 1:500 | Millipore-Sigma | SAB4200500 |
| Alpha-dystroglycan | Mouse | IgM | IIH6C4 | 1:100 | Developmental Studies Hybridoma Bank | IIH6 C4 |
| Glial fibrillary acid protein (GFAP) | Rat | Monoclonal IgG2a | 2.2B10 | 1:500 | Thermo-Fisher | 13-0300 |
| Vimentin | Mouse | IgG1 | V9 | 1:500 | Thermo-Fisher | MA5-11883 |
| Nestin | Mouse | Monoclonal IgG1 | Rat-401 | 1:100 | Millipore-Sigma | MAB353 |
| Beta-III-Tubulin (Tuj1) | Chicken | Polyclonal IgY | NA | 1:500 | Neuromics | CH23005 |
| Microtubule associated protein (MAP)-2 | Rabbit | Polyclonal | NA | 1:1000 | Thermo-Fisher | PA5-85756 |
| Tau | Chicken | Polyclonal | NA | 1:1000 | PhosphoSolutions | 1998-TAU |
| Postsynaptic density (PSD)-95 | Mouse | Monoclonal IgG2a | 6G6-1C9 | 1:500 | Thermo-Fisher | MA1-045 |
| Synaptophysin | Guinea Pig | Polyclonal | NA | 1:250 | Synaptic Systems | 101 004 |
| RBPMS | Guinea Pig | Polyclonal | NA | 1:500 | Millipore-Sigma | ABN1376 |
| **Secondary Antibodies** | | | | | | |
| **Antigen Target** | **Host Species** | **Fluorophore** |  | **Dilution** | **Company** | **Catalog Number** |
| Rabbit IgG | Goat | Alexa-488 |  | 1:500 | Thermo-Fisher | A11008 |
| Rabbit IgG | Goat | Alexa-568 |  | 1:500 | Thermo-Fisher | A11011 |
| Rabbit IgG | Goat | Alexa-647 |  | 1:500 | Thermo-Fisher | A32733 |
| Mouse IgG | Goat | Alexa-488 |  | 1:500 | Thermo-Fisher | A11029 |
| Mouse IgG | Goat | Alexa-568 |  | 1:500 | Thermo-Fisher | A11004 |
| Mouse IgG | Goat | Alexa-647 |  | 1:500 | Thermo-Fisher | A32728 |
| Mouse IgM | Goat | Alexa-488 |  | 1:500 | Thermo-Fisher | A21042 |
| Chicken IgY | Goat | Alexa-488 |  | 1:500 | Thermo-Fisher | A11039 |
| Rat IgG | Goat | Alexa-647 |  | 1:500 | Thermo-Fisher | A21247 |
| Goat IgG | Donkey | Alexa-555 |  | 1:500 | Thermo-Fisher | A32816 |
| Guinea Pig IgG | Goat | Alexa 488 |  | 1:500 | Thermo-Fisher | A11073 |

**Supplementary Table 1. Antibodies used for histology.**
