## Supplementary figures and images for "Structural engraftment and topographic spacing of transplanted human stem cell-derived retinal ganglion cells"

### Graphical Abstract

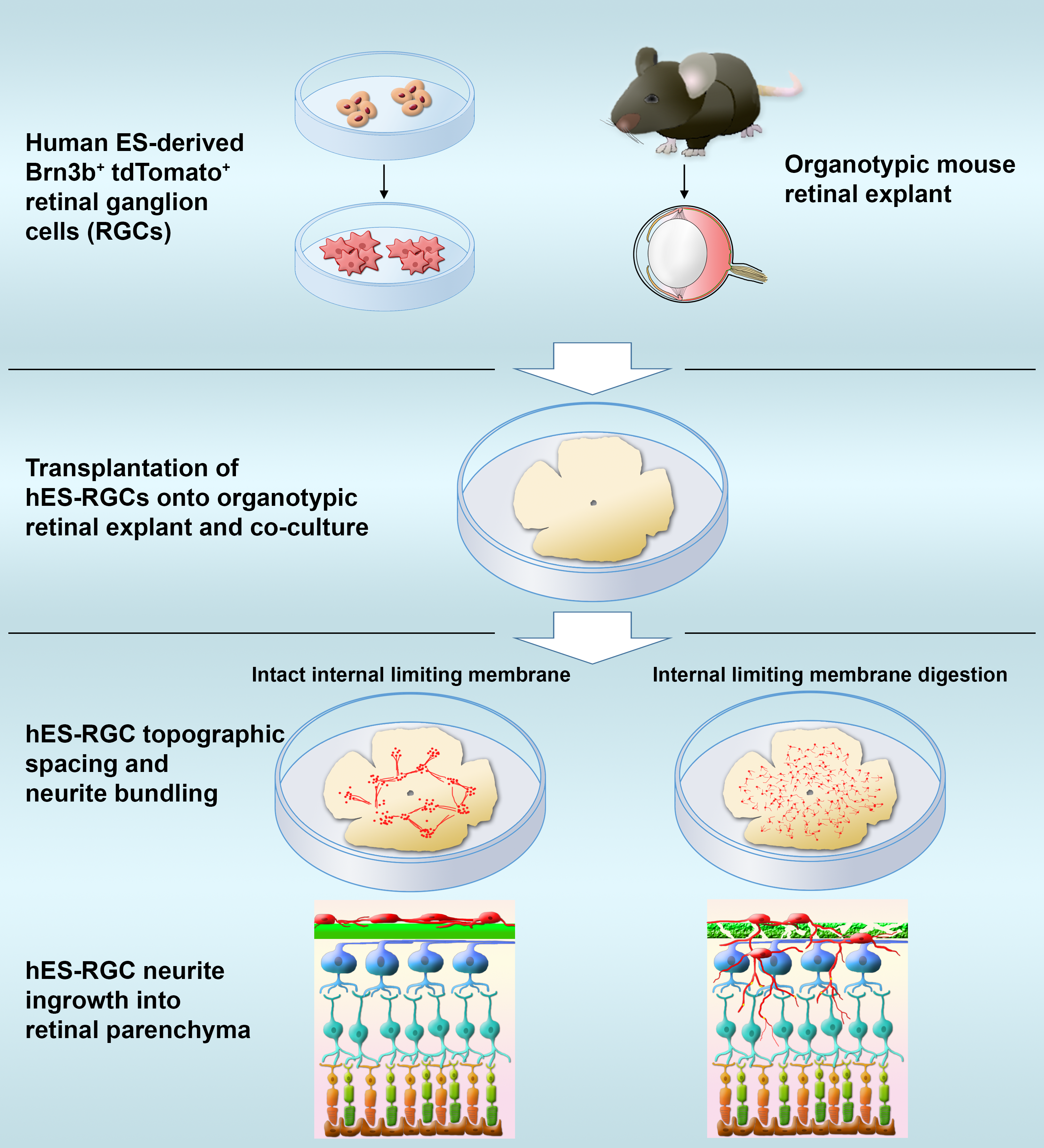

### Supplementary Figure 1

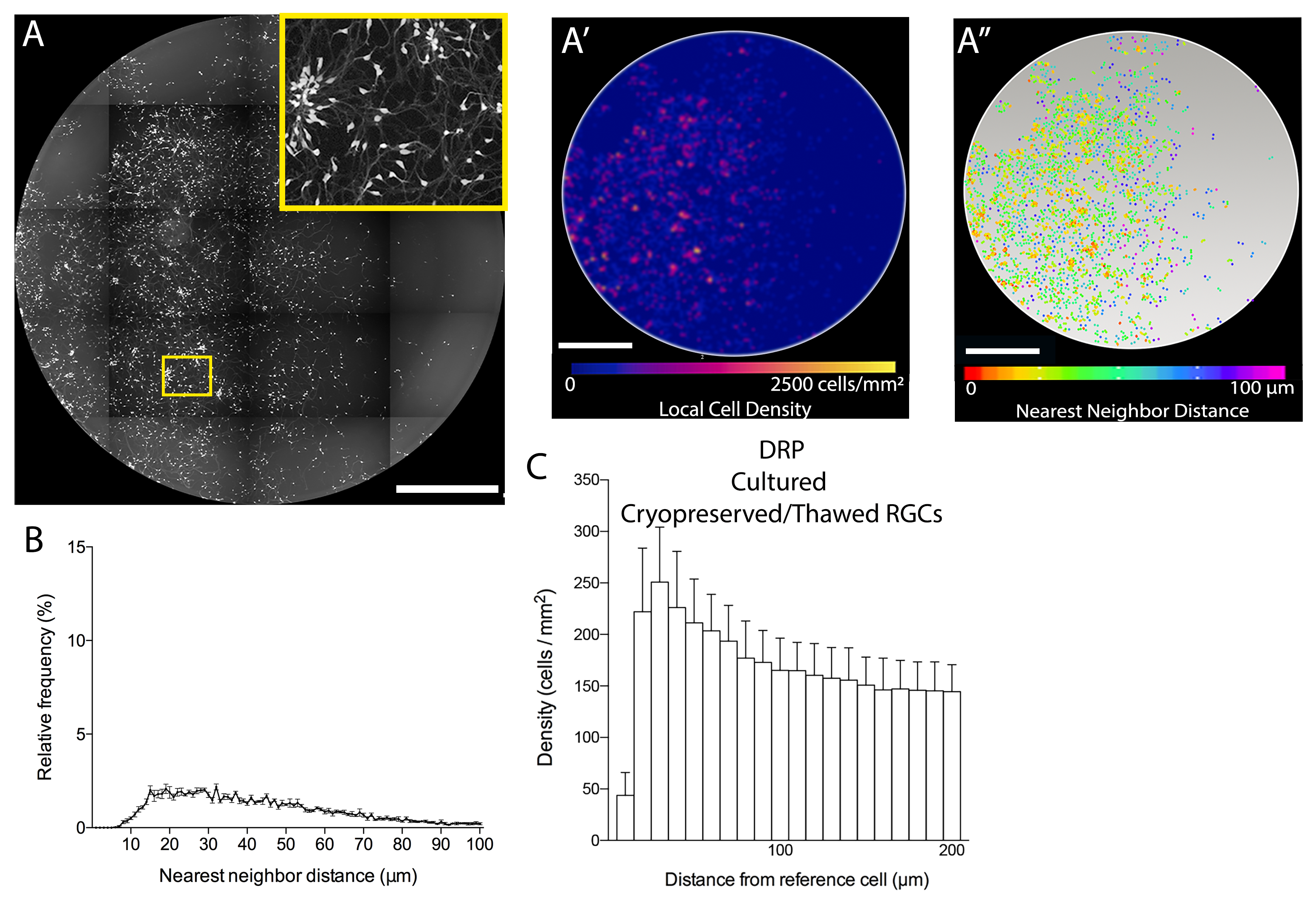

### Supplementary Figure 2

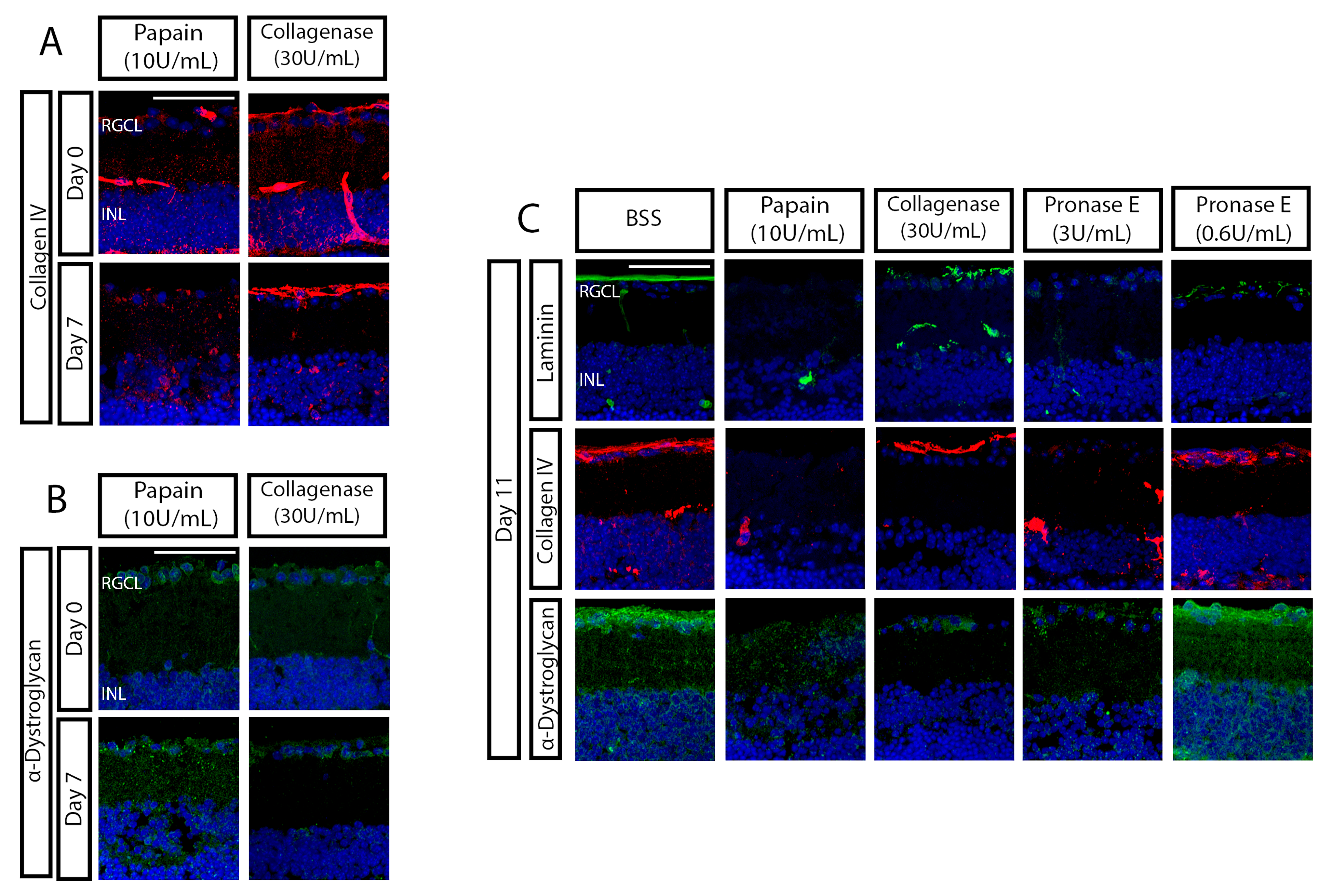

### Supplementary Figure 3

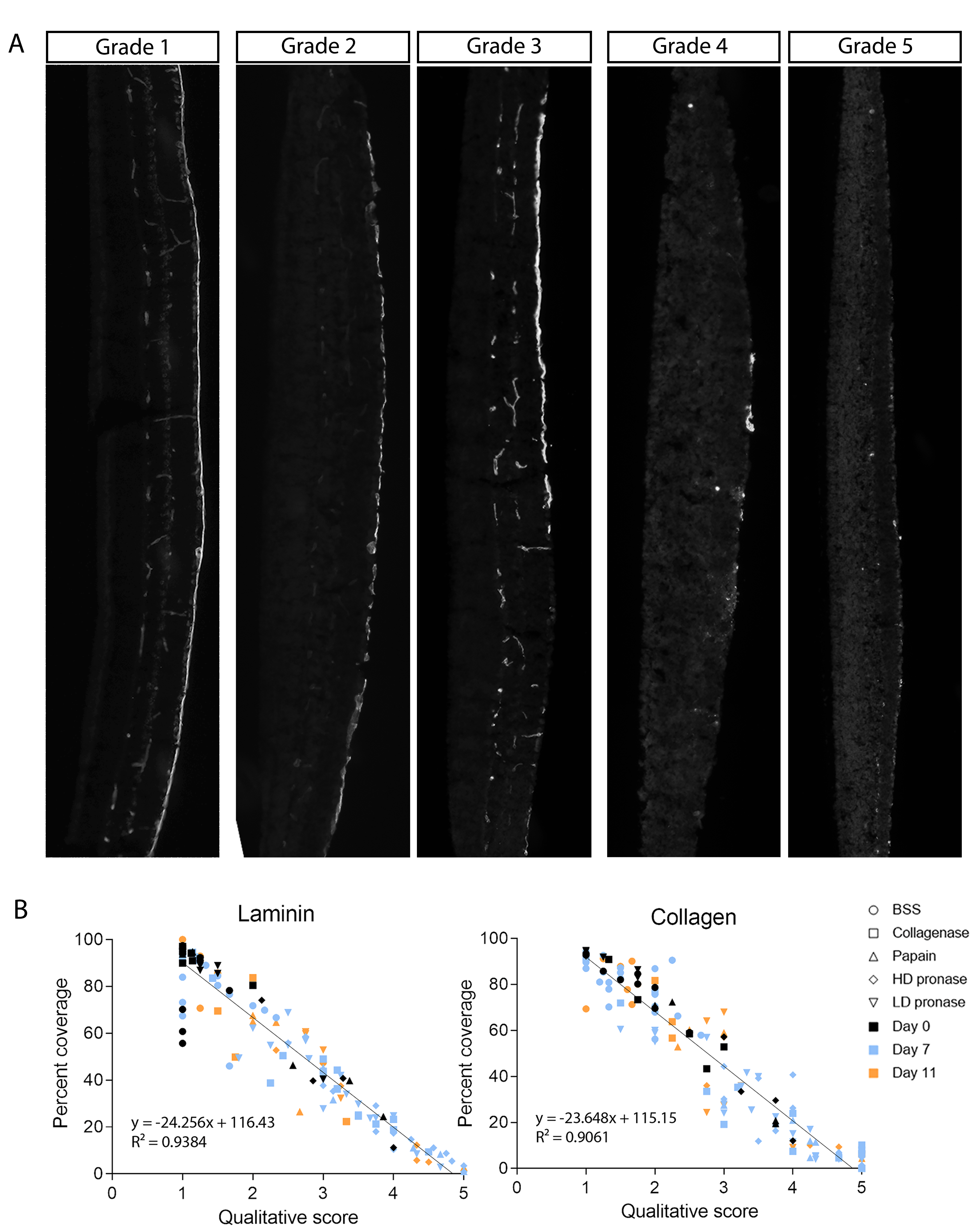

### Supplementary Figure 4

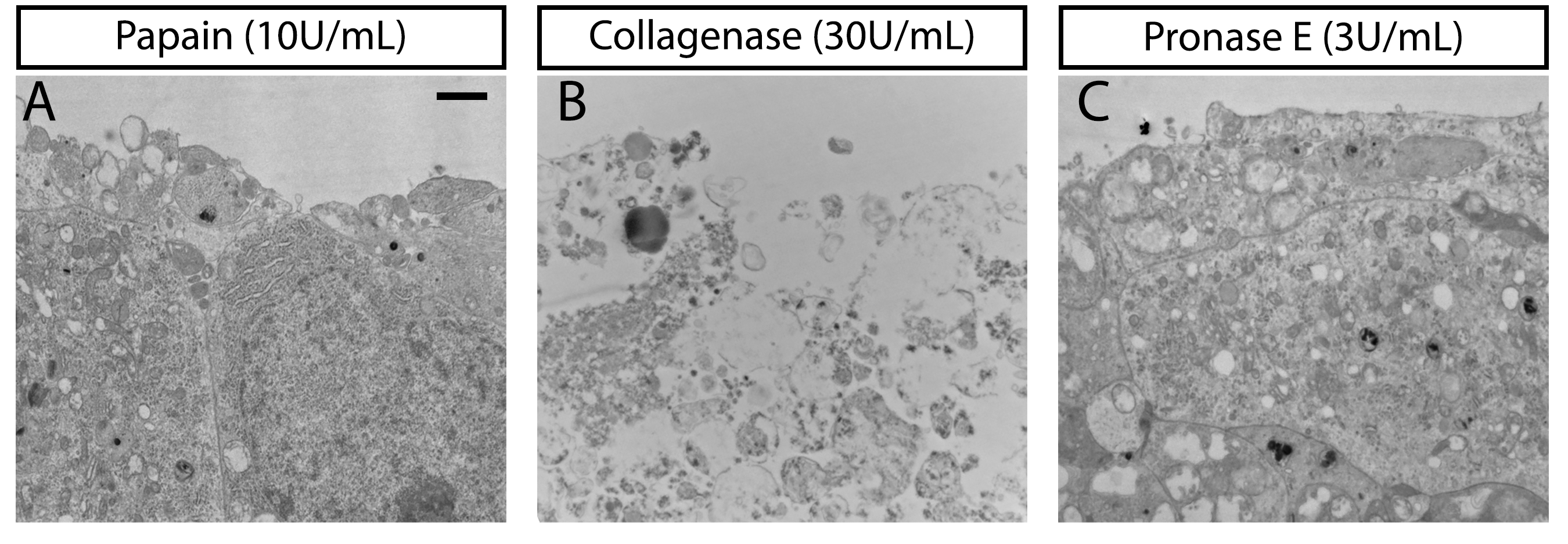

### Supplementary Figure 5

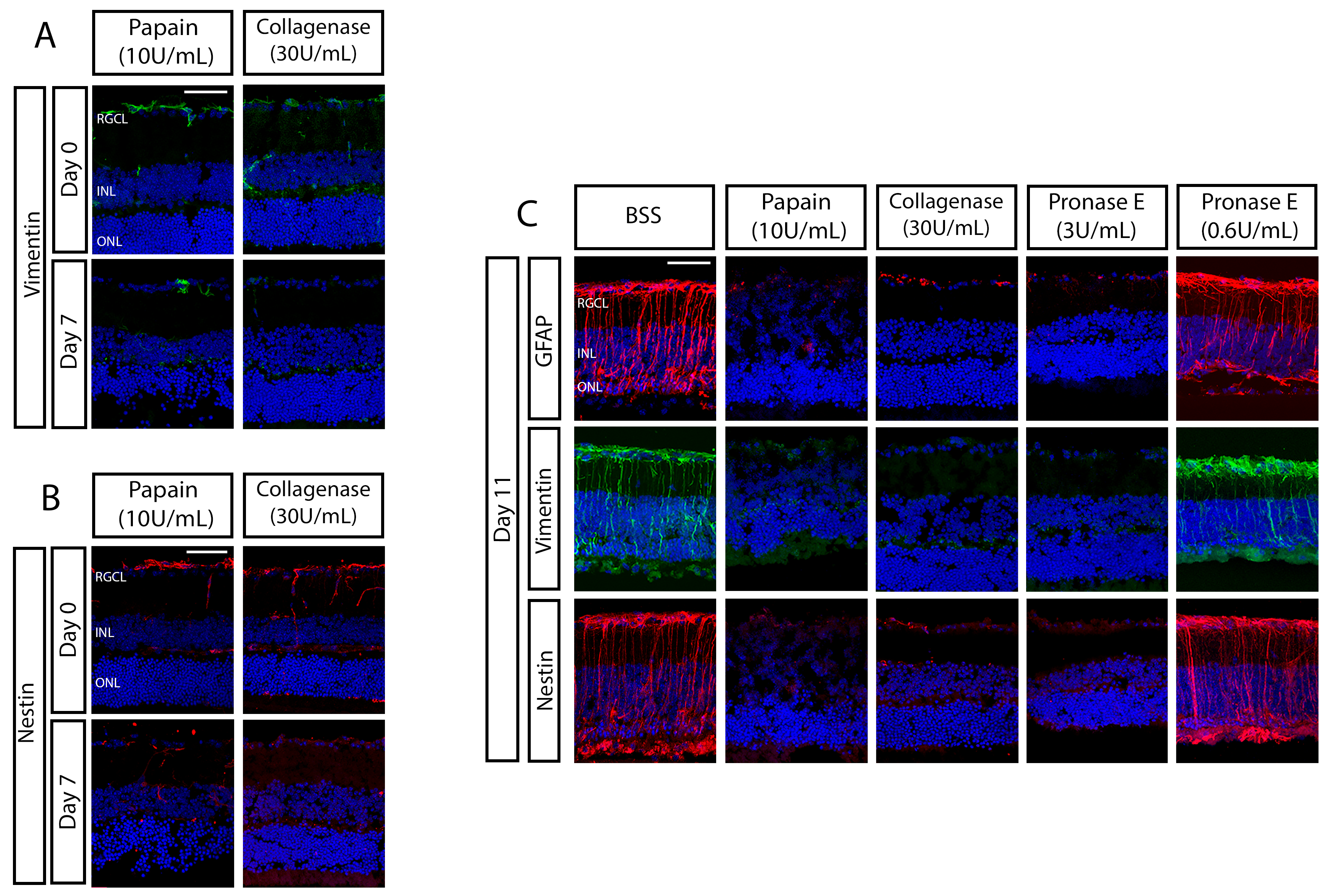

### Supplementary Figure 6

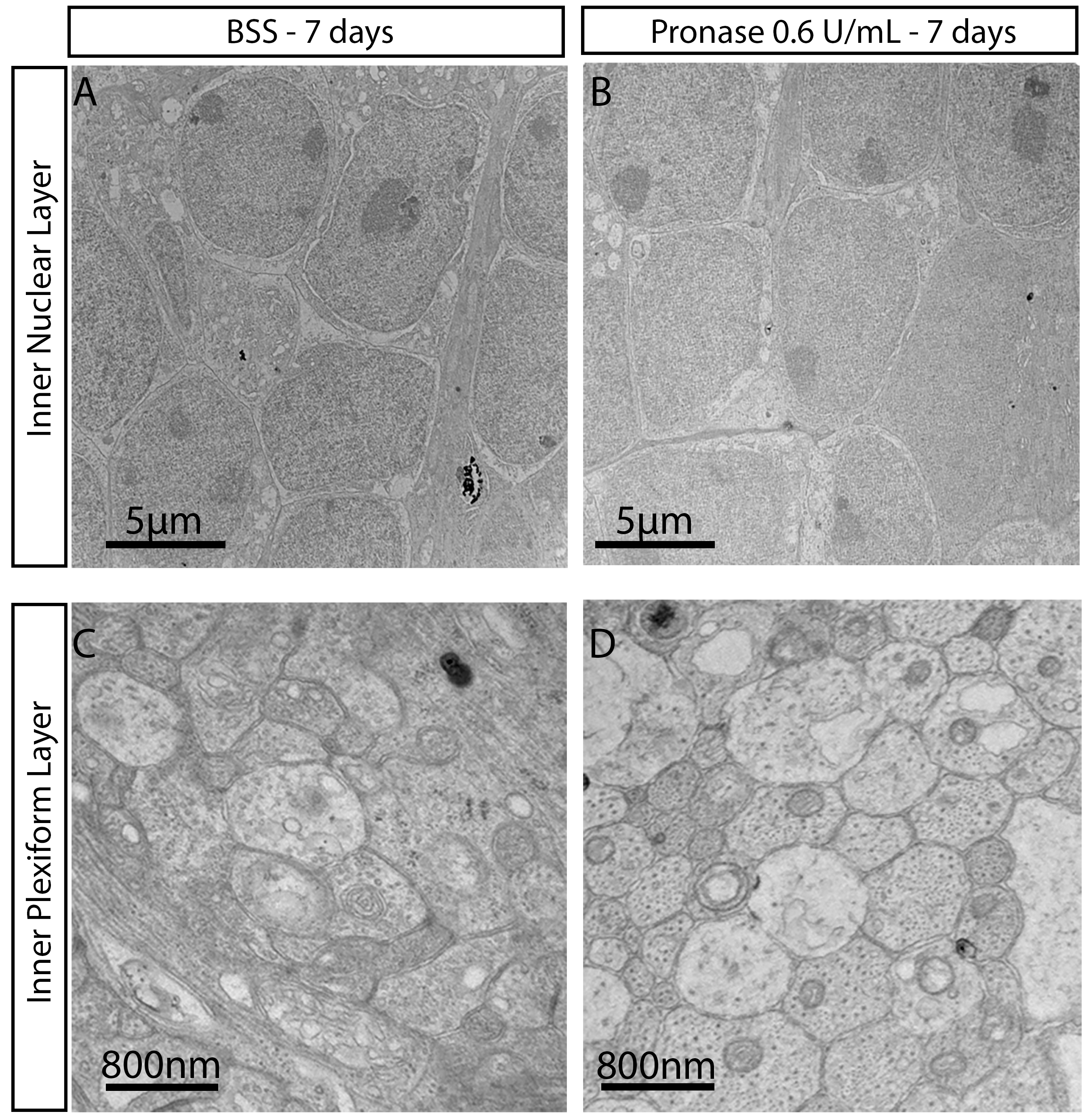

### Supplementary Figure 7

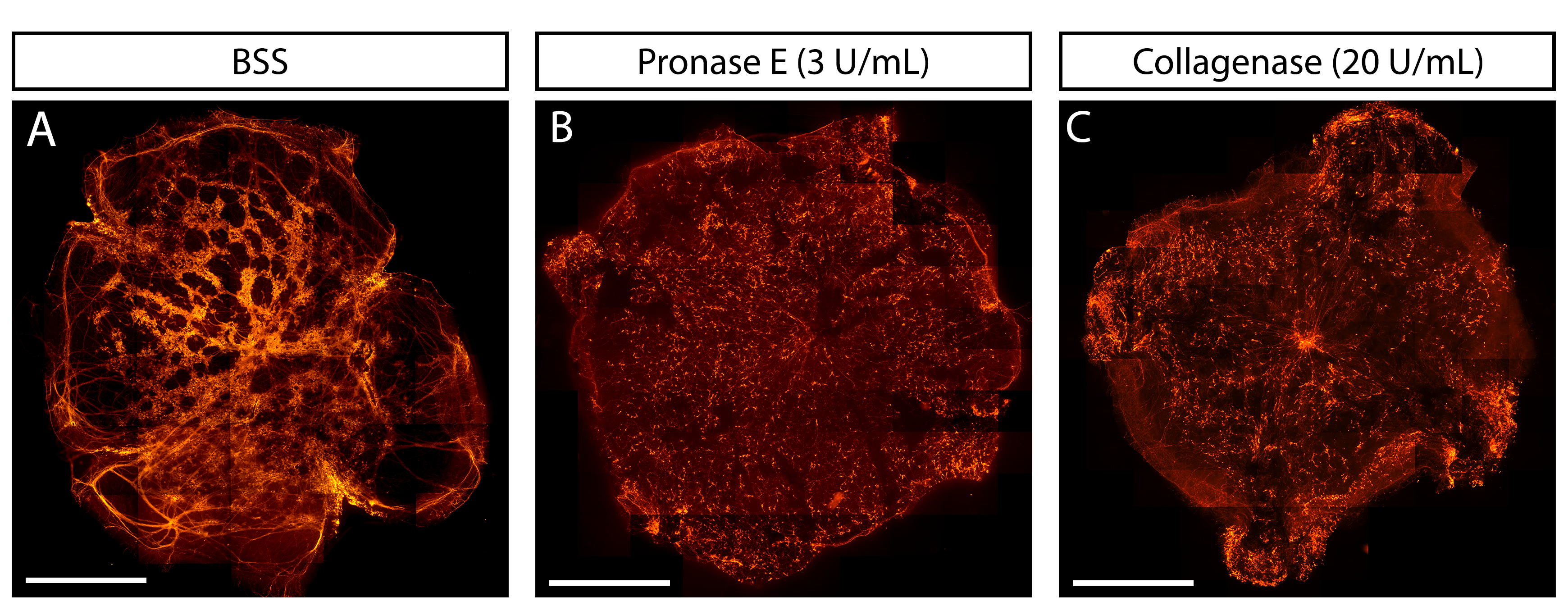

### Supplementary Figure 8

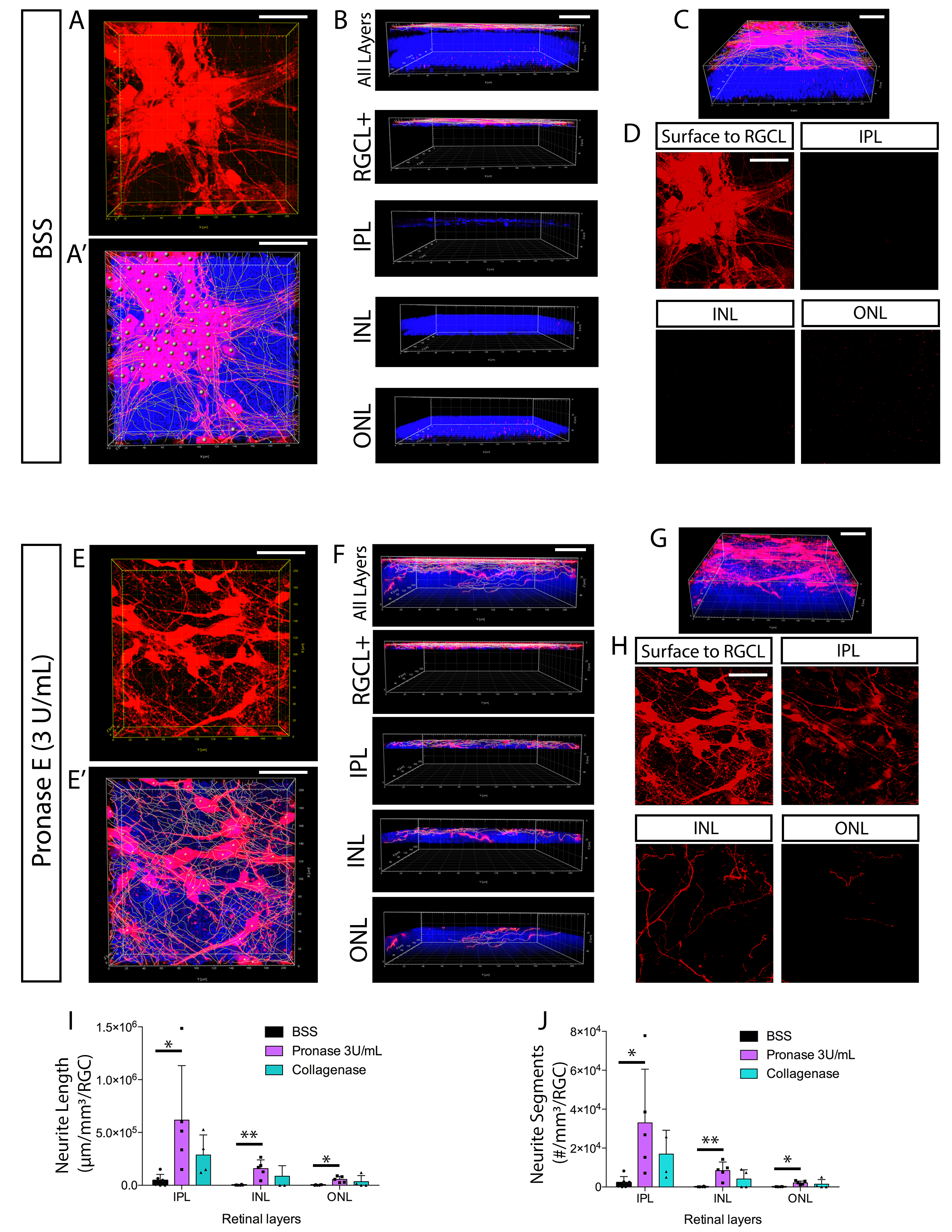
